# Naive primary neutrophils play a dual role in the tumor microenvironment

**DOI:** 10.1101/2023.09.15.557892

**Authors:** Kehinde Adebayo Babatunde, Rupsa Datta, Nathan W Hendrikse, Jose M Ayuso, Anna Huttenlocher, Melissa C. Skala, David J Beebe, Sheena C Kerr

## Abstract

The tumor microenvironment (TME) is characterized by a network of cancer cells, recruited immune cells and extracellular matrix (ECM) in a hypoxic microenvironment. However, the specific role of neutrophils during tumor development, and their interactions with other immune cells is still not well understood. Thus, there is a need to investigate the interaction between primary neutrophils and natural killer cells and the resulting effects on tumor development. Here we use both standard well plate culture and an under oil microfluidic (UOM) assay with an integrated extracellular cell matrix (ECM) bridge to elucidate how naive primary neutrophils respond to both patient derived tumor cells and tumor cell lines. Our data demonstrated that both patient derived head and neck squamous cell carcinoma (HNSCC) tumor cells and MDA-MB-231 breast cancer cells trigger cluster formation in neutrophils, and the swarm of neutrophils restricts tumor invasion through the generation of reactive oxygen species (ROS) and neutrophil extracellular trap (NETs) release within the neutrophil cluster. However, we also observed that the presence of neutrophils downregulates granzyme B in NK-92 cells and the resulting NETs can obstruct NK cells from penetrating the tumor mass *in vitro* suggesting a dual role for neutrophils in the TME. Further, using label-free optical metabolic imaging (OMI) we observed changes in the metabolic activities of primary neutrophils during the different swarming phases when challenged with tumor cells. Finally, our data demonstrates that neutrophils in direct contact, or in close proximity, with tumor cells exhibit greater metabolic activities (lower nicotinamide adenine dinucleotide phosphate (NAD(P)H) mean lifetime) compared to non-contact neutrophils.

## INTRODUCTION

The TME is a complex system, playing an important role in tumor development and progression. It is characterized by a network of cancer cells, with recruited immune cells such as neutrophils, and the extracellular matrix (ECM). Neutrophils are the most abundant leukocytes in the blood and are one of the first defenses against infection and tissue damage (*1*). The number of tumor-infiltrating neutrophils is a negative prognostic factor in several types of cancer (*2*). Several experimental and clinical studies have shown that increased neutrophil count and a higher neutrophil-to-lymphocyte ratio are associated with adverse outcomes in patients with cancer (*3*–*5*) thereby raising important questions on the impact of neutrophils in tumor development. In the context of tumor metastasis, the specific role of neutrophils in this process is still not clear. Thus, there is a need to elucidate the multifaceted roles played by neutrophils during tumor development.

While much immunological knowledge has been obtained from mouse studies, several components of the mouse immune system do not fully translate to the human immune system (*6*). While these animal models replicate microenvironmental complexity and physiological conditions, several aspects including species-specific differences and the limited ability to image immune cell-tumor interactions in real time make investigation of these interactions challenging in *in vivo* models.

Microfluidic technologies have overcome some of the challenges of traditional *in vitro* approaches based on Petri dishes by enabling the integration of 3D complex environment and cell-ECM interactions for modeling *in vivo*–like responses (*7*). Several microfluidic platforms for modeling immune cell-tumor cell interactions have been reported and they have contributed to understanding immune cell behavior in the TME (*8*, *9*). Previous studies have reported the use of microfluidic platforms to investigate how tumor cells trigger neutrophil extracellular traps in neutrophils (*9*) and the role of other immune cells such as T cells and NK cells during tumor development (*10*, *11*). However, most of these microfluidic platforms don’t accurately mimic the dissemination of tumor cells by invading into the ECM that can influence the dynamics of migration and the behavior of these immune cells in response to tumor cells.

Here, we use both standard well plate culture and an under oil microfluidic (UOM) assay (*12*, *13*) to conduct *ex vivo* studies of primary neutrophil functional responses when challenged with patient derived HNSCC tumor cells, a HNSCC tumor cell line and a breast cancer cell line (MDA-MB-231). These cell types were investigated because these cancer types are associated with increased neutrophil infiltration into the TME (*14*–*16*). Using these *in vitro* systems, we can visualize the dissemination of metastatic tumor cells (MDA-MB-231) by invading into the ECM, followed by neutrophil recruitment to the tumor site. By quantifying gene expression, cell death (via microscopy and flow cytometry) and the assessment of changes in the metabolic activity using label-free optical metabolic imaging (OMI) in neutrophils, we gained insights on neutrophil effector functions cooperating to either restrict or increase tumor invasion *in vitro*.

## Material and Methods

### UOM Device Construction

Briefly, a premium microscope slide (Fisher-finest, 3″ × 1″ × 1 mm; Thermo Fisher Scientific, 12-544-1) or chambered coverglass (1 well, no. 1.5 borosilicate glass, 0.13 to 0.17 mm thick; Thermo Fisher Scientific, 155360) was treated first with O2 plasma (Diener Electronic Femto, Plasma Surface Technology) at 60 W for 3 min and then moved to an oven for grafting of PDMS-Silane (1,3-dichlorotetramethylsiloxane; Gelest, SID3372.0) (about 10 μl per device) onto the glass surface by chemical vapor deposition (CVD) (*12*).

The PDMS-grafted slide was masked by a PDMS stamp and treated with O_2_ plasma at 60 W for 3 min. After surface patterning, the PDMS stamp was removed by tweezers and stored in a clean space for reuse. The glass slides were held in a plate [Nunc four-well tray, polystyrene (PS), non-treated sterile, Thermo Fisher Scientific, 267061] and overlaid with oil (silicone oil, 5 cSt; Sigma-Aldrich, 317667). The chambered cover-glass was directly overlaid with oil. For full details of the fabrication process, see Li et al. (*12*)

### ECM gel preparation

One-part rat tail collagen I (Corning, 10 mg/mL)) was neutralized with one-part neutrophil culture media (Roswell Park Memorial Institute (RPMI) 1640 Medium, Thermo Fisher Scientific,11875093) + 10% fetal bovine serum (Thermo Fisher Scientific, 10437010) and one-part 2X HEPES buffer. Channels were constructed by sweeping hanging droplets of collagen solution +/- cells across O2-plasma patterned areas. For full details of the channel preparation process, see Schrope et al. (*13*).

### Chemoattractant and cell loading

*f*MLP (Sigma-Aldrich), LTB4 (Cayman Chemical, Ann Arbor, Michigan, USA), IL-8 (Cayman Chemical, Ann Arbor, Michigan, USA) were diluted in RPMI to a working concentration of 100 nM. One microliter of the chemokine solution was pipetted into the chemoattractant spot (CAS) and one microliter of cell media suspension was then added to the cell loading spot (CLS) of the device. Finally, about 2□×□10^3^ primary neutrophils were pipetted into each CLS. Time-lapse images of neutrophil migration were captured using a fully automated Leica Super STED confocal microscope. The microscope is equipped with a bio chamber heated at 37◦C and 5% CO_2_.

### Cell culture

The NK-92 cell line was acquired from the American Type Culture Collection and cultured in X-VIVO 10 (Lonza, 04-380Q) supplemented with 20% fetal bovine serum (FBS; Lonza) and 0.02 mM folic acid (Sigma-Aldrich, F8758) dissolved in 1 N NaOH, 0.2 mM myo-inositol (Sigma-Aldrich, I7508), and IL-2 (100 U/ml; PeproTech, 200-02). Patient-derived HNSCC cells (IDC; 171881-019-R-J1-PDC, passages 7 to 12) were obtained from the National Cancer Institute Patient-Derived Models Repository (NCI PDMR). Patient-derived cells required specialized medium consisting of 1× Advanced Dulbecco’s modified Eagle’s medium/F12 (Gibco, 12634010) supplemented with 5% FBS, 1% penicillin/streptomycin, 0.178 mM adenine (Sigma-Aldrich, A8626-1G), 1.61 nM EGF Recombinant Human Protein (Invitrogen, PHG0313), 1.1 μM hydrocortisone (Sigma-Aldrich, H4001-1G), 2 mM l-glutamine (Invitrogen), and 0.01 mM Y-27632 dihydrochloride (Tocris). All tumor cell line was cultured in DMEM (30-2002, ATCC, Manassas, VA) supplemented with 100X non-essential amino acid (11140050, Gibco, ThermoFisher Scientific, Waltham, MA), 10% fetal bovine serum (FBS, 35-010-CV, Corning-cellgro, Manassas, VA) and 1% Penicillin-Streptomycin (Pen/Strep, 15,070,063, ThermoFisher Scientific, Waltham, MA).

### Isolation of human neutrophils

Neutrophils were isolated from peripheral human blood acquired by venipuncture from healthy donors under an approved IRB protocol at UW Madison. Isolation was performed within 1 h after the blood draw using the Easy sep human neutrophil negative isolation kit (STEMcell Technologies, Vancouver, Canada) following the manufacturer’s protocol. After isolation, neutrophils were stained with either cell tracker blue dye or calcein. Stained neutrophils were then suspended in RPMI 1640 media containing 10% FBS (Thermo Fisher Scientific) and penicillin/streptomycin.

### Neutrophil-mediated *in vitro* cytotoxicity assay

Neutrophils from healthy individuals were isolated for the cytotoxicity assay, as previously described. Cytotoxicity of tumor cells by human neutrophils was analyzed using microscopy and flow cytometry. Briefly, 2-3 × 10^3^ tumor cells were seeded into 96 well plates for 24 h. Human neutrophils from healthy individuals and NK-92 cells were added to achieve effector-to-target (E: T) cell ratios of 10:2:1 (neutrophil:NK:tumor). Cells were incubated at 37°C in a humidified 5% CO_2_ incubator for 24 h. Cytotoxicity was calculated by measuring the fluorescence intensity of calcein-AM, propidium iodide and caspase 3/7 dyes for live, non-caspase and caspase dependent cell death respectively. Stock solutions of calcein acetomethyl ester (CAM; 5 mg/ml) (Thermo Fisher Scientific, C3100MP), propidium iodide (PI; 2 mg/ml) (Thermo Fisher Scientific, P1304MP) and caspase 3/7 (Thermo Fisher Scientific) were prepared following the manufacturer instructions. Working solutions were prepared dissolving at 1:1000 and 1:500 respectively from the stock solutions in PBS while caspase 3/7 dye was directly added at 1:1000 in cell culture. After staining, cells were gently washed twice with PBS to remove dead cells and viable tumor cells were stained with the CAM solution. Images were acquired and evaluated using a Leica SP8 STED super-resolution confocal microscope.

### Cell isolation from tri culture

To selectively retrieve the neutrophils or NK-92 cells from tri-culture, two microliters of biotinylated anti-EpCAM (D20655, Thermo Fisher Scientific) and biotinylated CD56 (Thermo Fisher Scientific) were added to the well, and the sample was incubated at 4°C for 15 min. Ten microliters of Sera-Mag streptavidin coated beads (21152104011150, GE) were added, and the sample was incubated for another 10 min at 4°C. The Sera-Mag beads, with the tumor cells (EpCAM-positive) or NK cells (CD56 positive), were isolated using a magnet, leaving the neutrophils in suspension. Neutrophils were lysed using RLT plus buffer (Qiagen) for PCR analysis. Isolated NK cells with beads were also lysed using RLT buffer for PCR analysis.

### Flow cytometry

Following coculture, cells were harvested by using 0.25 % trypsin for 5-10 mins. For the different cell populations, we used cell tracker blue dye, vibrant red dye and cell tracker CMTPX (Thermo Fisher Scientific) to stain neutrophils, tumor cells and NK-92 cells respectively. Working solutions were prepared dissolving at 1:1000 the stock solutions in PBS for each dye. After detachment using trypsin and washing, cells were re suspended in FACS buffer (0.1% BSA in PBS and 2% FBS) containing 50µg/ml of DNase 1 (Sigma D-4513) to reduce clustering of cell samples. Fluorescence intensity was measured by flow cytometry. For cell surface marker analysis, cells were detached using 2mM EDTA for 15-20 mins. After detachment, cells were washed and stained with fluorochrome-conjugated CD56 antibody for NK-92 cells and CD11b antibody for primary neutrophils (Biolegend, San Diego, CA) for 30□min on ice. After staining cell was washed and suspended in FACS buffer (0.1% BSA in PBS, 2% FBS) at a concentration <1□×□10^7^ cells□ml^−1^. Fluorescence intensity was measured by flow cytometry (ThermoFisher Attune).

### Gating Strategy

Data was analyzed using FlowJo Software (Version 9). The gating strategy was as follows: (i) Cells of interest were obtained by gating on cell population based on size and granularity using (FSC vs SSC) to exclude debris (Supplementary Figure 4Bi). (ii) Single cells were identified by gating on cell of interest population by using forward scatter height (FSC-H) versus forward scatter area (FSC-A) density plot for double cells exclusion (Supplementary Figure 4Bii). (iii) Tumor cells were identified by gating on the single cell population negative for cell tracker blue (Neutrophils) (Supplementary Figure 4Biii) and/or cell tracker CMTPX (NK-92) (iv) Caspase 3/7 induced dead tumor cells were identified by gating on cells positive for caspase 3/7 dye and vibrant red dye (Supplementary Figure 4Biv).

### NETosis visualization and chemical inhibitors

To visualize NET formation, sytox green was added to the media at 500 nM (ThermoFisher Scientific). Quantification of NETs released within each cluster was performed by measuring the area covered by sytox green staining using the threshold and measure function in image J. 4-ABAH and MK-886 (Cayman Chemical, Ann Arbor, MI, USA), were dissolved in DMSO according to the manufacturer’s protocol. Each compound was re-suspended in cell culture media at a concentration of 20 µM for 4-ABAH, 400 nM for MK-886 pathway inhibitor. For MPO inhibition, neutrophils were incubated in 4-ABAH for 30 min and for LTB_4_ synthesis inhibition, neutrophils were incubated in MK-886 for 30 min before incubating with tumor cells.

### Tumor Spheroid Generation

Multicellular tumor spheroids were generated using the hanging drop method with methylcellulose (*17*). Briefly, 12 g of high viscosity methylcellulose (Sigma, M0512) were dissolved in one liter of RPMI1640, and centrifuged at 4000 g for 2.5 h. Only the clear supernatant was used in the next step. Cell suspension (10^5^ cells/ml) and this methylcellulose solution were mixed in a 4:1 v/v ratio, and 25 μl droplets (2×10^3^ cells) were placed on the lid of Petri dishes. Sterile water was placed on the bottom of the dishes to prevent evaporation from droplets, the lids were replaced, and the Petri dishes incubated at 37°C and 5% CO_2_. After 1 day, a single well-defined spheroid was generated per drop.

### Tumor spheroid neutrophil coculture assay

To evaluate the neutrophil recruitment ability of tumor spheroids in culture, non-tissue culture treated 96 well-plate were coated with 100 µg/ml fibronectin for at least 1 h. at 37°C. Approximately 15 spheroids were suspended in 200 µl fresh media. Spheroids were allowed to attach to the plate for about 4 h in the incubator at 37°C/5% CO_2_. 1×10^5^ primed neutrophils were seeded to each well in 200 µl of 1% serum media. We imaged the samples for 24 hours to confirm the migration of neutrophils toward tumor spheroid.

### Image Processing and Data Analysis

Microscopy images were processed in Fiji ImageJ. The area covered by neutrophils and NK-92 cells were measured by setting an automatic threshold to the images, followed by the measure and/or analyze particle functions in Image J. The set automatic thresholds are in either Triangle or Huang modes to measure the area of cell tracker blue (NK-92 cells) or calcein (neutrophils) positive objects in the image. The graphs were plotted using excel and/or GraphPad Prism. The number of immune cells around a tumor spheroid was measured automatically using the Trackmate module in ImageJ. Using the Trackmate function on Image J, the “estimated cell sizes” was set at 10 μm and “intensity thresholds” to 2 for the immune cells. Parameters were selected based on the cell size and fluorescent intensity of cells in Image J. Kymograph and migrating speed of the immune cells were determined by using the kymograph and manual tracking plugin respectively in Image J.

### Optical metabolic imaging (OMI)

A custom-built inverted Ultima Multiphoton Imaging System (Bruker Fluorescence Microscopy, Middleton, WI) was used to acquire fluorescence intensity and lifetime images. The equipment consists of an ultrafast laser (Insight DS+, Spectra Physics), an inverted microscope (Nikon, Eclipse Ti), and a 40× water immersion 1.15 NA objective (Plan Apo, Nikon). NAD(P)H and FAD images were obtained for the same field of view, sequentially. FAD fluorescence was isolated using an emission bandpass filter of 550/100 nm and excitation wavelength of 890 nm. NAD(P)H fluorescence was isolated using an emission bandpass filter of 440/80 nm and an excitation wavelength of 750 nm. A dichroic mirror at 720 nm was used to remove excitation photons from emission. Fluorescence lifetime data were collected using time-correlated single-photon counting electronics (SPC-150, Becker and Hickl) and a GaAsP photomultiplier tube (H7422P-40, Hamamatsu). Imaging was performed using Prairie View Software (Bruker). Images (256 × 256 pixels) with an optical zoom of 2, were obtained using a pixel dwell time of 4.8 μs over 60-s total integration time. The instrument response function (IRF) was collected each day by recording the second harmonic generation (SHG) of urea crystals (Sigma-Aldrich) excited at 890 nm.

For imaging, neutrophils monoculture and co-culture with primary tumor cells were seeded in the ratio of 10:1 (neutrophil to tumor cells) on glass bottom well plates. Throughout imaging, the samples were maintained at 37°C and 5% CO2. For metabolic inhibition, cells were treated with 2 deoxy-glucose (2-DG) (100mM) and carbonyl cyanide-4-(trifluoromethoxy)-phenylhydrazone (FCCP) (660nM).

NAD(P)H lifetime images were analyzed using SPCImage software (Becker & Hickl). To provide more robust calculations of the fluorescence lifetimes, background pixels with a low intensity were excluded by applying a threshold. The photon counts in each decay were increased by applying a a 3×3 bin consisting of 9 surrounding pixels. The fluorescence lifetime decay curves at each pixel were fit to a two-component exponential decay model by iterative re-convolution with the IRF at each pixel, *I*(*t*) = α1\**e*^(−t/τ1)^ + α2\**e*^(−t/τ2)^ + *C*, where *I*(*t*) represents the fluorescence intensity at time *t* after the laser excitation pulse, α accounts for the fractional contribution from each component, *C* represents the background light, and τ is the fluorescence lifetime of each component. Since NAD(P)H can exist in two conformational states, bound or unbound to enzymes, a two-component model was used (*18*). The short and long lifetime components reflect the free or unbound and bound conformations for NAD(P)H. The mean lifetime (τ_m_) was calculated using τ_m_ = α_1_τ_1_+ α_2_τ_2_for NAD(P)H. The optical redox ratio [NAD(P)H/NAD(P)H+FAD] was determined from the NAD(P)H and FAD by integrating the photons detected at each pixel in the image to calculate the total intensity (*19*). Single cell segmentation was performed using Cellpose 2.0 (model: cyto2), a deep learning-based segmentation method (*20*). Masks generated by Cellpose were manually checked and edited using Napari (CZI). Values for NA(P)DH τ_m_, NAD(P)H intensity, FAD intensity, and the optical redox ratio were averaged for all pixels within each cell mask. Neutrophils were divided into “contact” and “noncontact” subsets based on their spatial proximity to the tumor in the same coculture such that neutrophils approximately < 30µm from tumor cells were assigned to the “contact” group.

### Reverse transcription quantitative polymerase chain reaction

To study the molecular changes in neutrophils and NK-92 cells *in vitro*, the expression of neutrophil related genes was analyzed by RT-qPCR. Briefly, mRNA was isolated using the iQ SYBR Green Supermix kit (Bio-Rad). mRNA was reverse transcribed to complementary DNA (cDNA) using the iScript PreAMP cDNA Synthesis Kit (Bio-Rad) using primers from Integrated DNA Technologies (Supplementary Table 5). cDNA was analyzed by RT-qPCR using selected genes. cDNA was analyzed by RT-qPCR using iTaq Universal SYBR Green Supermix (Bio-Rad) or Roche LightCycler master mix according to the manufacturer’s protocols in Roche’s LightCycler 480 II. Gene expression was normalized using the ΔΔ^Ct^ method. Each experiment was run in triplicate.

### Statistical analysis

Statistical significance of the differences between multiple data groups were tested using two-way Analysis of Variance (ANOVA) in GraphPad Prism (GraphPad Software, version 9.0). Within ANOVA, significance between two sets of data was further analyzed using two-tailed t-tests. Non-paired Student’s t-test was performed for determining statistical significance between two conditions. All error bars indicate standard errors of means (SEM). ns, not significant (*p* ≥ 0.05); \**p*< 0.05; \*\**p* < 0.01; \*\*\**p* < 0.001. The number of replicates ranged from n = 3 to n = 5 for each experimental condition. All data are presented as means from 3 independent experiments. Data analysis for the OMI data was performed in Python, and graphs for all the figures were plotted in Python using the Seaborn data visualization library. Statistical tests were performed on jamovi (*21*). Data were compared using either Student’s t test or one-way analysis of variance (ANOVA) with a or Tukey’s post hoc test for multiple comparisons, and P values have been indicated as follows: \**p*< 0.05; \*\**p* < 0.01; \*\*\**p* < 0.001.

## RESULTS

### Quantifying coordinated migration in neutrophil in real time

To investigate the dynamics of neutrophil coordinated migration toward tumor cells, we first investigated the ability of tumor cells to recruit neutrophils *in vitro*. The UOM platform is based on the principle of exclusive liquid repellency (ELR) under silicone oil to create a 3D microenvironment, that can include an extracellular matrix (type 1 collagen) coating (*22*). While this UOM platform has previously been used to demonstrate neutrophil migration during mechanical constriction (*13*) here, we demonstrate how coordinated migration in neutrophils restricts tumor invasion in real time. The UOM migration design is an open, fluid, walled platform that is made up of two spots each with a diameter of 100µm; the cell loading spot (CLS), and the chemoattractant spot (CAS) which are connected by an ECM bridge channel of 500µm in length (Figure 1A). The UOM assay used here mimics tumor migration via the extracellular matrix and thus allows the study of the interaction between primary neutrophils and tumor cells in real time. To investigate neutrophil tumor cell interaction, we first established the functionality of the UOM assay by validating the chemotactic response of neutrophils to several known and established chemo-attractants. We chose N-formyl-Met-Leu-Phe (*f*MLP), a pro-resolution chemoattractant secreted by bacteria after infection, an intermediary chemoattractant leukotriene B_4_, (LTB_4_), and a pro inflammatory chemokine Interleukin (IL-8) (*23*).

**FIGURE 1:**
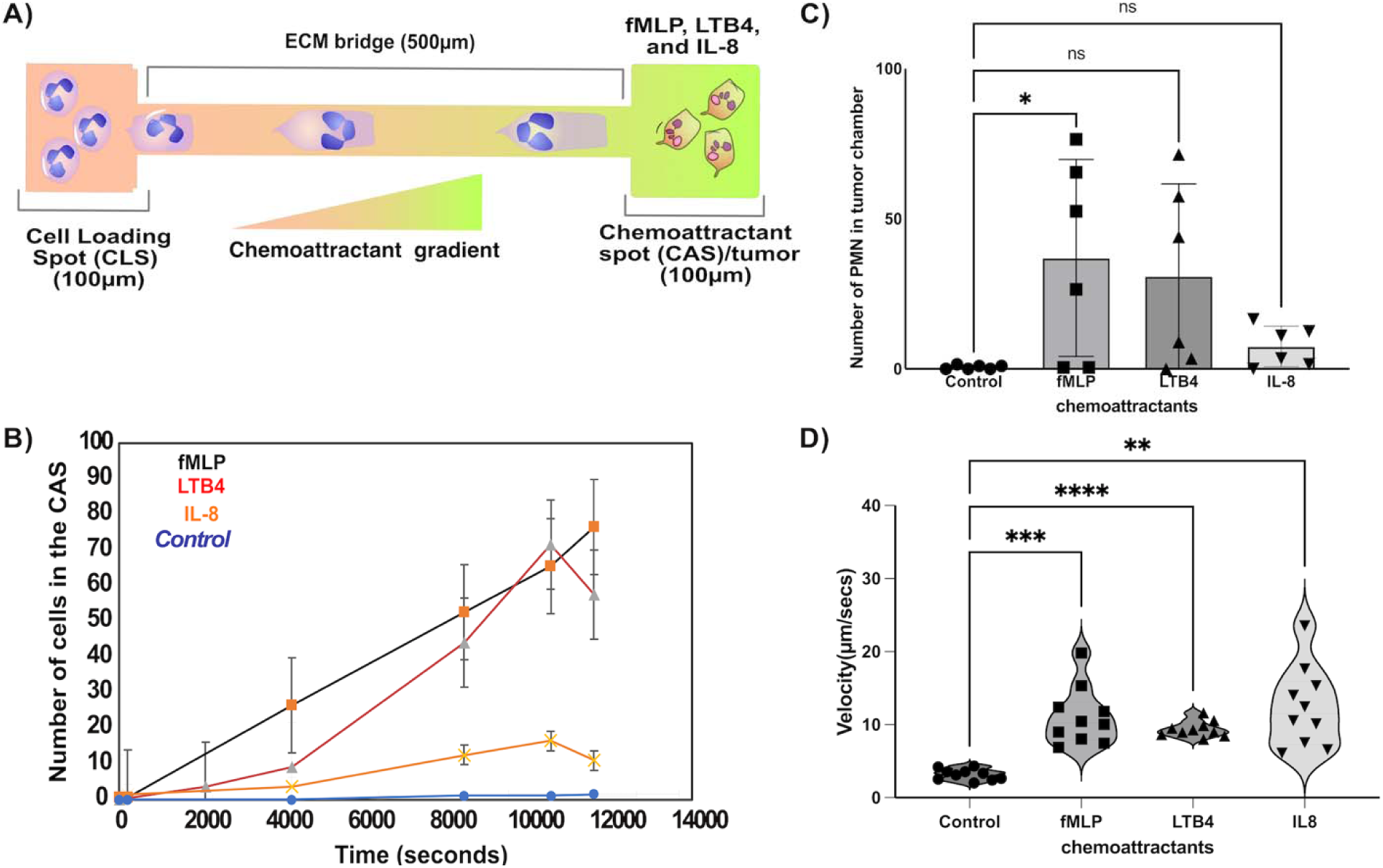
Quantification of neutrophil migration. **(A)** Schematic image illustration showing neutrophils in the cell loading spot (CLS) (100µm) migrating toward the chemoattractant gradient via the extracellular cell matrix (ECM) bridge (500 µm) to the chemoattractant spot (CAS) (100µm) of the UOM device. **(B)** Graph showing migratory dynamic of neutrophils in different chemo-attractant gradients migrating out of the CLS toward the ECM bridge and into the CAS. *f*MLP (Black line), LTB_4_ (red line), IL-8 (orange line) and control (blue line). (C) Graph showing the number of neutrophils that migrated out of the CLS toward the ECM bridge and into the CAS at 3-hour time point. **(D)** Graph showing the velocity of neutrophil migrating in the different chemoattractant gradients. Each dot represents a neutrophil. Comparison of velocity of migration in control to each chemo-attractants were performed by one-way ANOVA; \*\*\**P*□<□0.001 in control-*f*MLP, \*\*\*\**P*□<□0.0001 in control-LTB4 and \*\**P*□<□0.01 control-IL-8. Schematic illustration was designed by Affinity Designer Software (version 1.10.6.) https://affinity.serif.com/en-us/designer/.

To perform the assay, we delivered chemoattractant into the CAS to create a chemokine gradient between the two ELR spots. Neutrophils loaded into the CLS migrated directionally through the ECM bridge into the CAS. Tracks of migrating neutrophil trajectories were generated by analyzing time-lapse images using Image J software (Supplementary Figure 1A-D). The outcome of this chemotactic response was classified as persistent migration into the CAS. Neutrophils demonstrated significant migration towards the fMLP gradient, a trend towards increased migration towards LTB4, with lower levels of migration towards IL-8 (Figure 1B-C). In the *f*MLP gradient the total number of neutrophils that migrated into the CAS was increased two-fold compared to the IL-8 gradient (Figure 1C) as previously described (*24*). The average speed of migrating neutrophils in control (PMN without chemo-attractant), *f*MLP, LTB_4_ and IL-8 were 3.17µm/min, 11.1µm/min, 9.4µm/min and 10.8µm/min respectively (Figure 1D). Taken together, our data further confirm that neutrophils exhibit increased directional migration to a *f*MLP gradient compared to an IL-8 gradient (*24*) and that the UOM assay can be used to study coordinated migration in neutrophils during tumor challenge *in vitro*.

### Naive primary neutrophils form cluster around a tumor monolayer and restrict metastatic tumor cells in the UOM

Having validated that we can study neutrophil migration dynamics using the UOM assay, we next sought to interrogate the direct interactions between neutrophils and tumor cells in real time. We performed initial characterization in a well plate before moving into the UOM assay. We seeded primary patient derived HNSCC tumor cells in a monolayer for 24 h prior to co-culture with primary neutrophils (Figure 2A). Neutrophil and tumor cell interactions were examined via fluorescence microscopy after 24 hours in co-culture. We observed that the HNSCC tumor monolayer induced neutrophil clustering (Figure 2A-B). The observed neutrophil aggregation around tumor cells is similar to that described against zymosan targets (*25*, *26*) and is suggestive of a swarming process mediated by neutrophil-released leukotriene B (LTB_4_) (*27*). To verify the mechanism involved in the formation of neutrophil clusters around the tumor cells, we tested the role of LTB_4_ release during neutrophil aggregation by disrupting LTB_4_ synthesis with a MK886 pathway inhibitor (Figure 2C), which reduced neutrophil swarming. These data suggested that this clustering is LTB_4_ dependent. The area of clustering formed in the control condition is significantly larger than that of the MK-886 treated neutrophils (∼□400,000 μm^2^ vs. 100,000 μm^2^, for control vs. MK-886 treated neutrophils) (Figure 2D).

**FIGURE 2:**
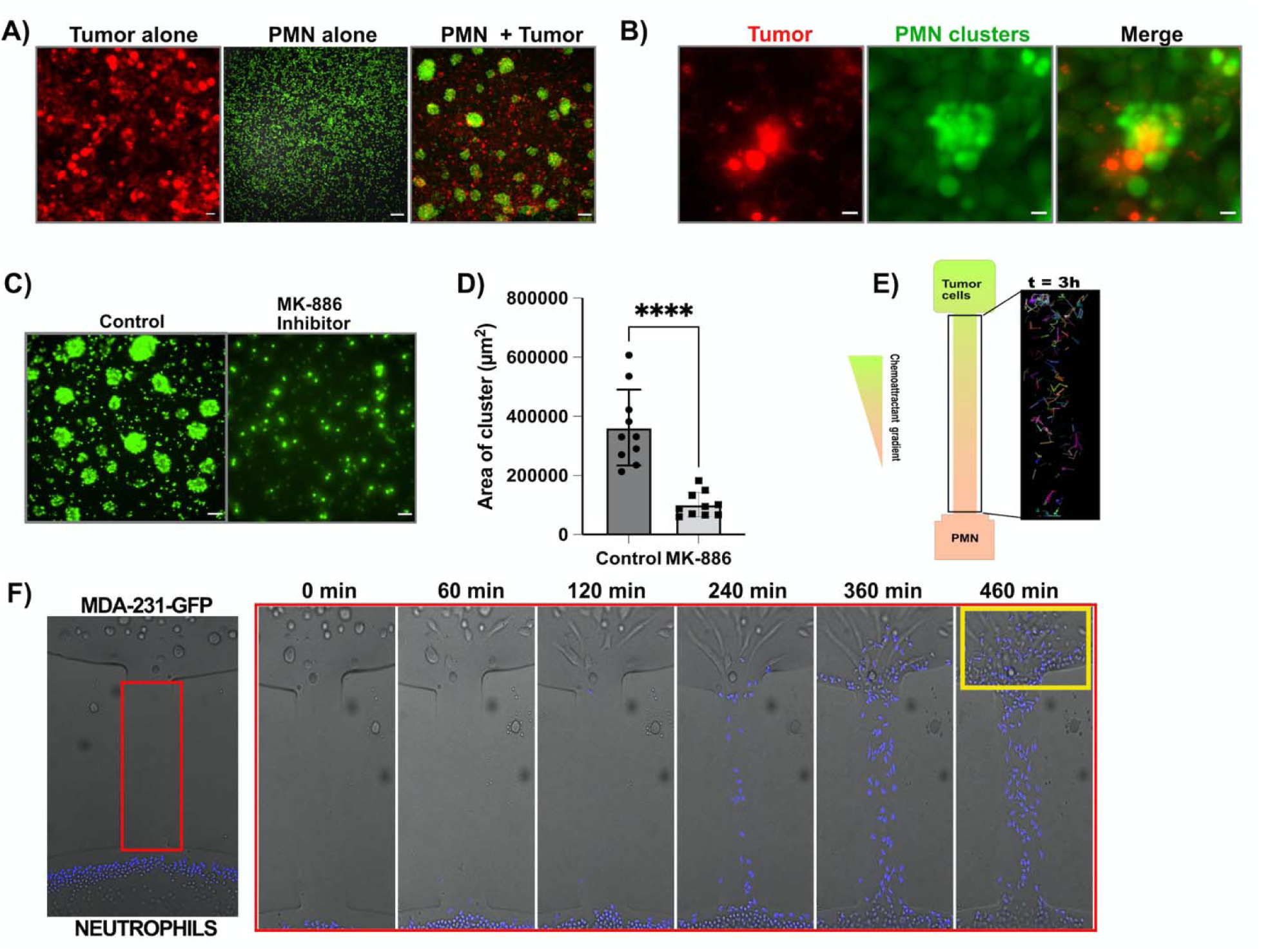
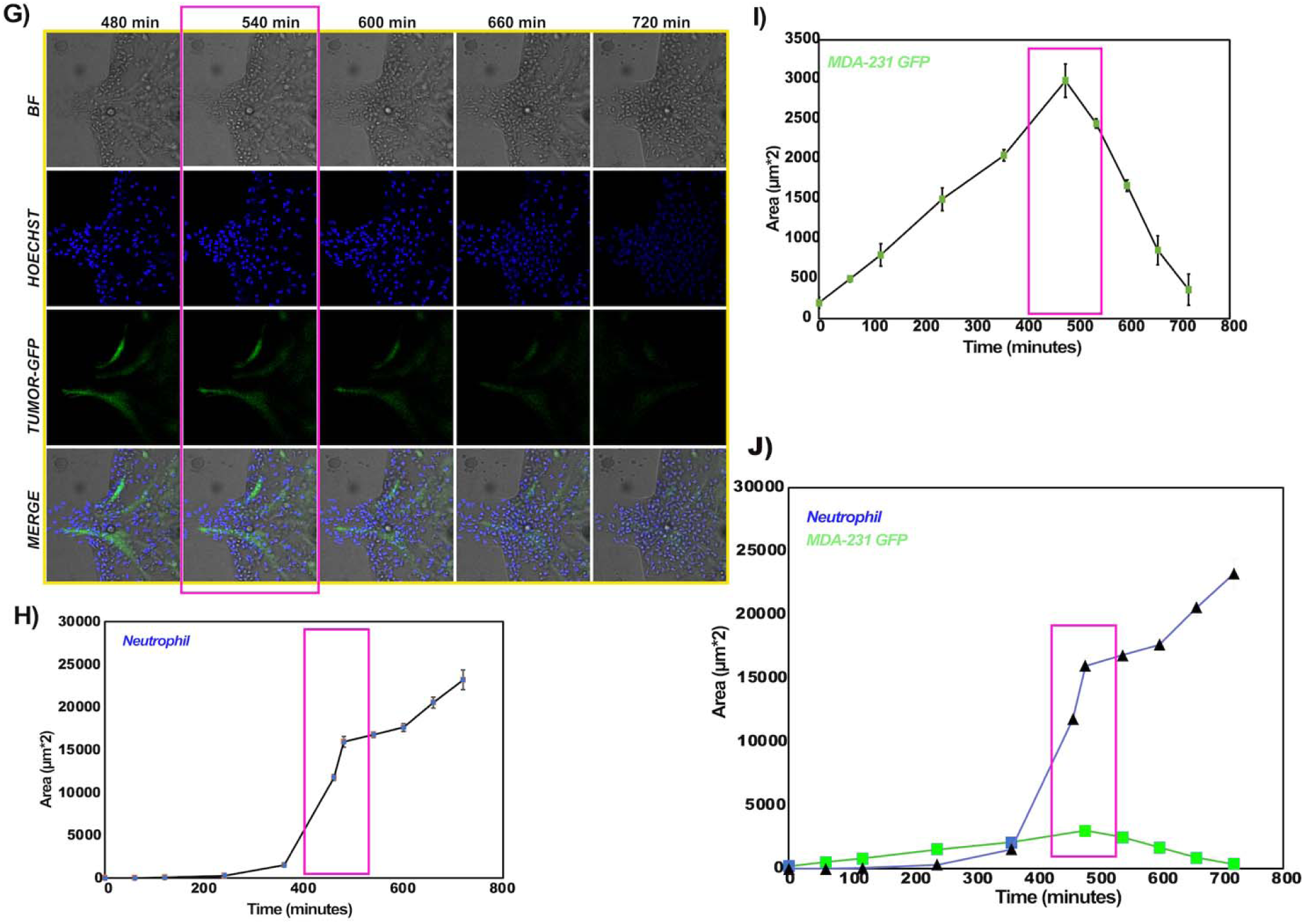
Primary neutrophils restrict metastatic tumor cells in the UOM. **(A)** Showing images of neutrophils clustering around the tumor monolayer after 24 hours. Tumor cells and neutrophils were stained with vibrant red dye and calcein dye respectively. **(B)** Showing magnified images of neutrophils clustering around the tumor monolayer. **(C)** Primary neutrophils form smaller clusters in the presence of LTB_4_ pathway inhibitors (MK-886). **(D)** Quantification of cluster formation in primary neutrophils and MK-886 treated neutrophils. Comparison between the area covered by control and MK-886 were performed by unpaired t test; \*\*\*\**P*□<□0.0001. Figure shows representative experiments of N□=□20 clusters. **(E)** Schematic illustration showing migratory tracks of neutrophils migrating from CLS toward tumor cells in the CAS. **(F)** Sequential time images showing clustering of primary neutrophils (blue) in response to tumor cells (Bright field) at 0 min, 60 min, 120 min, 240 min, 360min and 460 min. **(G)** Sequential images showing cluster formation in primary neutrophils (blue) around MDA-MB-231^GFP^ (green) at 480 min, 540 min, 600 min, 660 min and 720 min. Graph showing the migratory dynamics of **(H)** neutrophil, **(I)** metastatic MDA-MB-231 GFP and **(J)** neutrophil and MDA-MB-231 GFP (merged) by measuring the area (µ m*2) covered over time (minutes).

Having observed neutrophil clusters around tumor cells in our well plate culture, we next sought to understand the migratory dynamics in neutrophils during tumor challenge using the UOM assay. To do this, we used MDA-MB-231 tumor cells because they are a highly aggressive tumor cell line which exhibited a high migration and invasive capability (*28*). Thus, we first loaded MDA-MB-231 tumor cells in the CLS and quantified baseline levels of tumor migration along the ECM bridge (Video 1A). Next, we loaded MDA-MB-231 tumor cells in the CAS and monitored neutrophil chemotactic migration from the CLS toward the tumor cells using time lapse microscopy. PMN tracks were recorded using Image J software (Figure 2E).

Tumor cell migration starts when the morphology of the cell changes from a small rounded shape to an elongated spindle like shape after 60 minutes of incubation in the UOM. At 1-2□h, tumor cells start to migrate toward the ECM bridge. In response to the migrating tumor cells, we first observed individual neutrophils migrating toward the tumor chamber (*29*)(Figure 2F), which was immediately followed by robust neutrophil recruitment to the tumor chamber via the ECM bridge (Video 2). The observed neutrophil swarm formed in the tumor chamber is similar to the migratory pattern observed in neutrophil swarms in infection (*27*, *30*). More specifically, several phases have been previously reported during the swarming process (*25*, *31*). Our data shows that during the first 120 minutes, neutrophils migrate directionally toward the tumor cells along the ECM bridge prior to cluster formation (Figure 2F & Video 2). After the first set of migrated neutrophils interact with the tumor cells, the number of migrating neutrophils toward the tumor increases rapidly to consolidate the cluster size (240-600 min- Figure 2F & Video 2). Neutrophil clusters reach their peak size 660–720 min later, after which the cluster size remains stable (Figure 2J & Video 2).

To better visualize the migration of the tumor cells along the ECM bridge, we employ the use of a metastatic cancer cell line strain expressing GFP (MDA-MB-231*^GFP^*) (Figure 2G). The swarm size gradually increases at ∼8h in the tumor chamber (Figure 2G & H). The increase in the neutrophil cluster size gradually reduces the tumor migration and metastasis from the CAS to the ECM bridge *in vitro* (Figure 2I & J) (Video 2) compared to tumor alone (Video 1A). Taken together, these data demonstrate that primary tumor cells can trigger recruitment and cluster formation in neutrophils, and the observed clustering is LTB_4_ dependent.

### Tumor cells trigger NETs release in naive primary neutrophil *in vitro*

Following the observation of cluster formation around tumor cells, we sought to investigate if NETs were released in these neutrophil clusters. To do this, we used immunofluorescence staining to detect specific NET-associated proteins such as myeloperoxidase (MPO) and citrullinated histone (cit-H3) positive neutrophils in our co-culture (Figure 3A-B). Our immunofluorescent staining shows co-localization of MPO^+^ and citrullinated histone H3^+^ neutrophils in the group co-cultured with tumor cells (Figure 3B-insert 2 & C-insert 1), however, we observed less citrullinated histone H3^+^ neutrophils (Figure 3D). While we could visualize NETs in vitro, to investigate if there was evidence of NET production in primary tumor tissue, we carried out immunofluorescent staining to detect NET markers MPO and cit-H3 in human breast cancer tissue (Figure 3C). We also observed co-localization of MPO^+^ and citrullinated histone H3^+^ neutrophils in our primary tissue (Figure 3C-insert 1). Using the UOM assay, next we showed that NETs are released during the neutrophil-tumor interactions in real time. To do this, we monitored the release of NETs along the ECM bridge. NETs release was quantified by examining the appearance of Sytox green staining along the ECM bridge. We found that at the point of interaction between neutrophils and tumor cells along the ECM bridge, extracellular DNA was released by neutrophils as detected by the appearance of sytox green staining (Figure 3E) (Video 3A & 3B). The area of sytox green starts to gradually increase at 4h after the direct contact between neutrophils and migrating tumor cells (Figure 3F).

**FIGURE 3:**
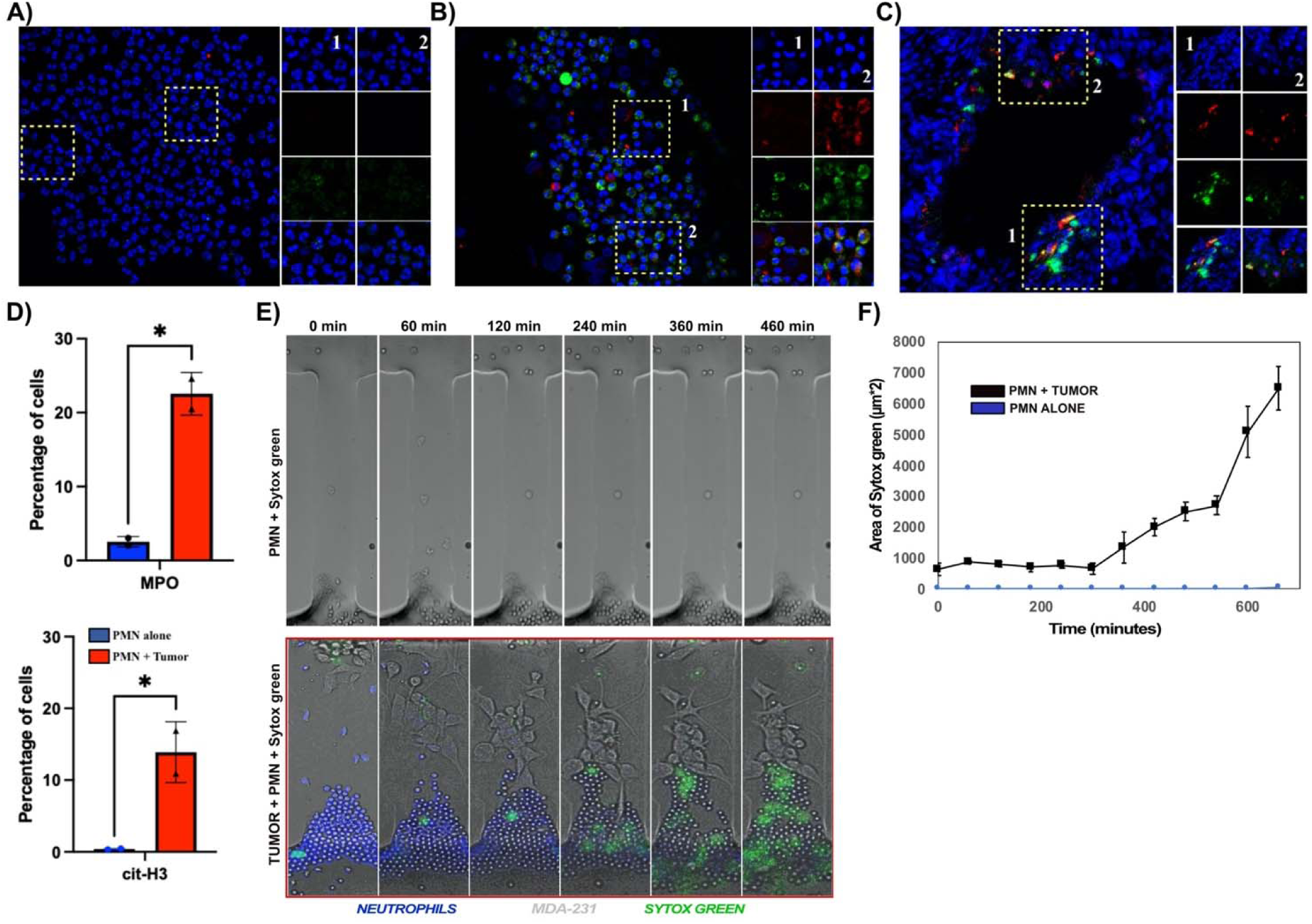
Tumor cells trigger NETs release in primary neutrophils *in vitro*. Immunofluorescence staining showing specific NETs protein-citrullinated histone H3^+^ and myeloperoxidase (MPO^+^) staining in **(A)** Neutrophils alone **(B)** Neutrophils co-cultured with tumor cells and **(C)** Human breast cancer tissue. Neutrophils (Blue), MPO (green) and Cit-H3 (Red). **(D)** Graph showing the percentage of cells expressing MPO and Cit-H3 in A and B. Quantification of extracellular DNA released by neutrophils during tumor interaction was done by measuring the **(E)** Sequential images of the appearance of sytox green dye along the ECM bridge in the UOM at 0 min, 60 min, 120 min, 240 min, 360min and 460 min. **(F)** Dynamics of the appearance of sytox green dye along the ECM bridge in PMN alone (blue line) and PMN + Tumor (black line) was recorded by measuring the area covered by sytox green (µ m*2) over time (minutes). Comparison between the area covered by MPO and cit-H3 in PMN alone and PMN + tumor was performed by unpaired t test; \**P*□<□0.05.

### Patient derived tumor cells trigger a change in metabolic activity in naive neutrophils

Having shown that tumor cells trigger a swarming response in neutrophils in the UOM assay, we hypothesized that the metabolic activity of neutrophils could be altered at the different phases of swarming [*scouting stage (0-120min), amplification (6-7h) and stabilization stage (20-24h)*]. Thus, we sought to investigate the metabolic activity of neutrophils in monoculture and co-culture by label free optical metabolic imaging (OMI), which quantifies the mean lifetime of reduced form of nicotinamide adenine dinucleotide phosphate [NAD(P)H] and the optical redox ratio (*19*) in single cells (Figure 4A) during the different stages of neutrophil swarming. We found that compared to the neutrophils in monoculture, the neutrophils in coculture exhibited a significantly lower mean NAD(P)H lifetime (NAD(P)H τ_m_) and a higher fraction of free NAD(P)H (α_1_). These findings at 0h were observed for neutrophils isolated from 2 individual blood donors (Supplementary Figure 2A, Supplementary Table 2). Moreover, co-culture neutrophils had a significantly higher NAD(P)H τ_m_ and a lower NAD(P)H α_1_ during the *scouting phase* (0-0.5 hour) compared to co-culture neutrophils during the *amplification phase* (6-7 hours) (Figure 4B, Supplementary Table 1). This could indicate a shift of metabolism to glycolysis in neutrophils in the presence of tumors (*32*). Finally, at 20-24 hours during the (s*tabilization phase*), there was no significant difference between neutrophils in the mono- and co-culture conditions (Supplementary Figure 2A, Supplementary Table 2). The comparable metabolic characteristics in both the experimental conditions is expected by 24 hours since neutrophils undergo either NETosis (in the co-culture condition) or apoptosis (in the monoculture condition) over time.

**FIGURE 4:**
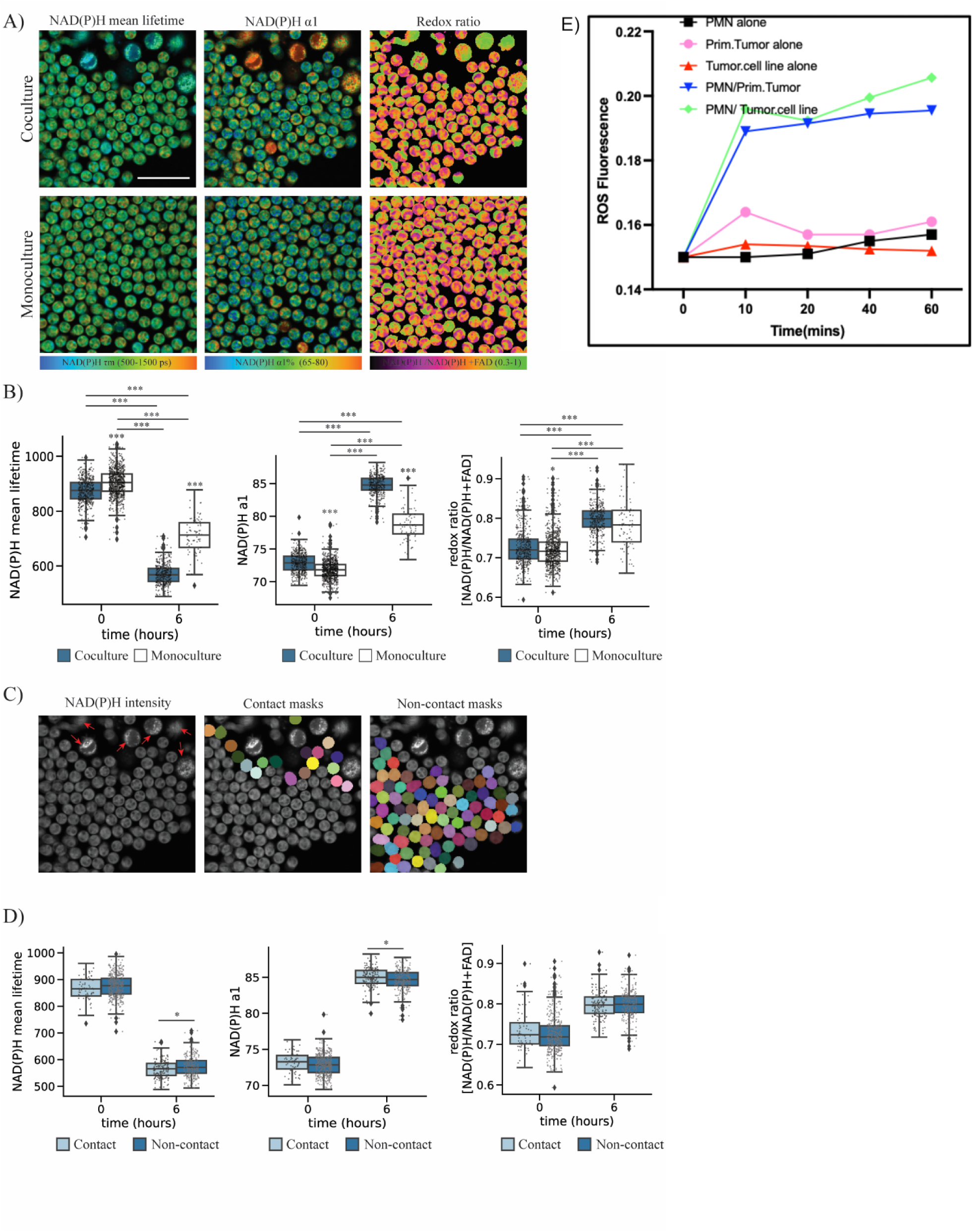
Autofluorescence imaging of neutrophil tumor interaction. **(A)** Representative images of NAD(P)H τ_m_, NAD(P)H α1% and optical redox ratio [NAD(P)H / (NAD(P)H + FAD)] of neutrophils cultured with and without primary HNSCC tumor cells. Scale bar = 50µm. **(B)** Quantification of NAD(P)H τ_m_, NAD(P)H α1% and optical redox ratios of neutrophils cultured either in monoculture or coculture with tumor cells before (0-1 hour) ‘0’ and after (6-7 hour) ‘6’ swarming response. Optical imaging data is analyzed for n = 9 images per condition from the 3 independent dishes. The data are computed at the single-cell level and the number of cells segmented per condition is presented in table S1. Statistical significance was accessed using one-way ANOVA with Tukey’s post hoc test (***P < 0.001; *P <0.5). Data is presented as medians and error bars are the 95% confidence interval. **(C)** Left -representative NAD(P)H intensity image (gray scale) of co-culture with tumor cells indicated with red arrows. NAD(P)H intensity image (gray scale) overlaid with contact (middle) and non-contact (right) neutrophil masks in color. The cells that were spatially less than approximately 30 µm from tumor cells are grouped as ‘contact’ while the rest of the cells were assigned to ‘non-contact’ group. **(D)** Quantification of NAD(P)H τm, NAD(P)H α1% and optical redox ratios of contact and non-contact neutrophils in coculture with tumor for ‘0’ (0-1 hours) and ‘6’ (6-7 hours) Optical imaging data is analyzed for n = 9 images per condition from the 3 independent dishes. The data are computed at the single-cell level and the number of cells segmented per condition is presented in table S3. Statistical significance was assessed using Student’s T-test (*P <0.5). Data is presented as medians and error bars are the 95% confidence interval. **(E)** Graph shows ROS production in neutrophil co-cultured with primary tumor cells (blue line) and tumor cell line (green line) via the measurement of released intracellular H_2_O_2_.

Importantly, within the co-culture condition, we observed that neutrophils in direct contact or close to the tumor cells during the *amplification phase* (6-7 hours) exhibited higher glycolytic activity (lower NAD(P)H τ_m_ and higher NAD(P)H α_1_) compared to neutrophils not in contact with tumor cells (Figure 4C-D, Supplementary Table 3). Together, these findings suggest that neutrophils are more likely to exhibit their effector functions when close to tumor cells by increasing their glycolytic rate. To explore the metabolic pathways neutrophils, employ during tumor challenge, we inhibited a glycolysis pathway and the pentose phosphate pathway (PPP) in neutrophils using 2 deoxy-glucose (2-DG) and carbonyl cyanide-4-(trifluoromethoxy)- phenylhydrazone (FCCP), respectively. 2-DG is a glucose analog and acts as a competitive inhibitor of glycolysis at the step of phosphorylation of glucose by hexokinase (*33*) and FCCP is an ionophore that disrupts ATP synthase and uncouples oxidative phosphorylation thus depleting mitochondrial membrane potential (*34*). We observed that unlike the untreated group, 2-DG and FCCP treated neutrophils in co-culture did not display significant differences compared to neutrophils alone (Supplementary Figure 2B). This suggests that both glycolytic/PPP pathways and mitochondrial metabolism might play a role in the observed metabolic shift in neutrophils under tumor challenge. The optical redox ratio did not show any significant differences in the experiments suggesting the redox state might not be affected by the presence of tumor.

To further confirm the metabolic changes in neutrophils cocultured with tumor, we co-incubated freshly isolated primary neutrophils with serum opsonized tumor cells for 1 hour and measured ROS production compared to primary neutrophil alone (control). To quantify extracellular ROS production in neutrophils, we used the H_2_O_2_□specific probe Amplex Red to measure extracellular H_2_O_2_ production in response to tumor exposure in neutrophils. Detectable H_2_O_2_ was observed within 10-20 min of exposure to both primary tumor cells and tumor cell lines. However, our data show significant differences in extracellular H_2_O_2_ production between control and tumor-neutrophils after 60 mins, as ROS production was significantly increased in tumor-neutrophils co-culture (Figure 4E). Taken together these data suggests that tumor challenge induces metabolic changes in neutrophils such that they are activated to produce ROS which could be both NOX and mitochondria derived. Moreover, the observation supports interaction between the cell types such that a direct contact or close proximity to tumor cells drives neutrophil activation.

### Neutrophil trigger caspase 3/7 pathway in tumor cell using the UOM

Tumor associated neutrophils (TANs) have been demonstrated to directly exert tumor death *in vitro* (*35*). However, it’s still unclear if isolated neutrophils from healthy donors could exert similar cytotoxicity against tumor cells *in vitro*. Thus, we sought to investigate whether neutrophils isolated from healthy donors can induce tumor cell death. For the killing experiments, we carried out parallel experiments with the same neutrophil donors on patient-derived tumor cells and a tumor cell line to minimize the issue of donor variability in our study. Using the UOM, we loaded MDA-MB-231 tumor cells (non-GFP) and neutrophils into the CAS and CLS respectively. Prior to loading, we pre-treated tumor cells with caspase 3/7 Cell Event dye for 30 mins and washed off excess staining prior to loading. Neutrophils loaded into the CLS migrated directionally through the ECM bridge toward the tumor cells (Figure 5A). Caspase 3/7 induction was monitored by examining the appearance of caspase 3/7 event dye (apoptosis fluorescent dye) staining in the tumor chamber. We observed caspase 3/7 fluorescent signal (green) in the CAS (Figure 5A) (Video 4A & 4B). Using the well plate culture killing assay, we added caspase 3/7 dye and killing was confirmed via fluorescent microscopy (Figure 5B) and flow cytometry (Figure 5C). In our standard well plate killing assay, we labelled cells with caspase 3/7 dye to identify apoptotic cells in addition to labelling tumor cells and neutrophils with vibrant red dye and cell tracker blue dye respectively to permit their identification in the mixed co-culture. This allowed us to specifically identify that we were reporting tumor cell death. We found that primary neutrophils increased the percentage of dead tumor cells from ∼ 10 % in tumor monoculture to ∼ 25% in the neutrophil-tumor coculture (Figure 5Bi-ii).

**FIGURE 5:**
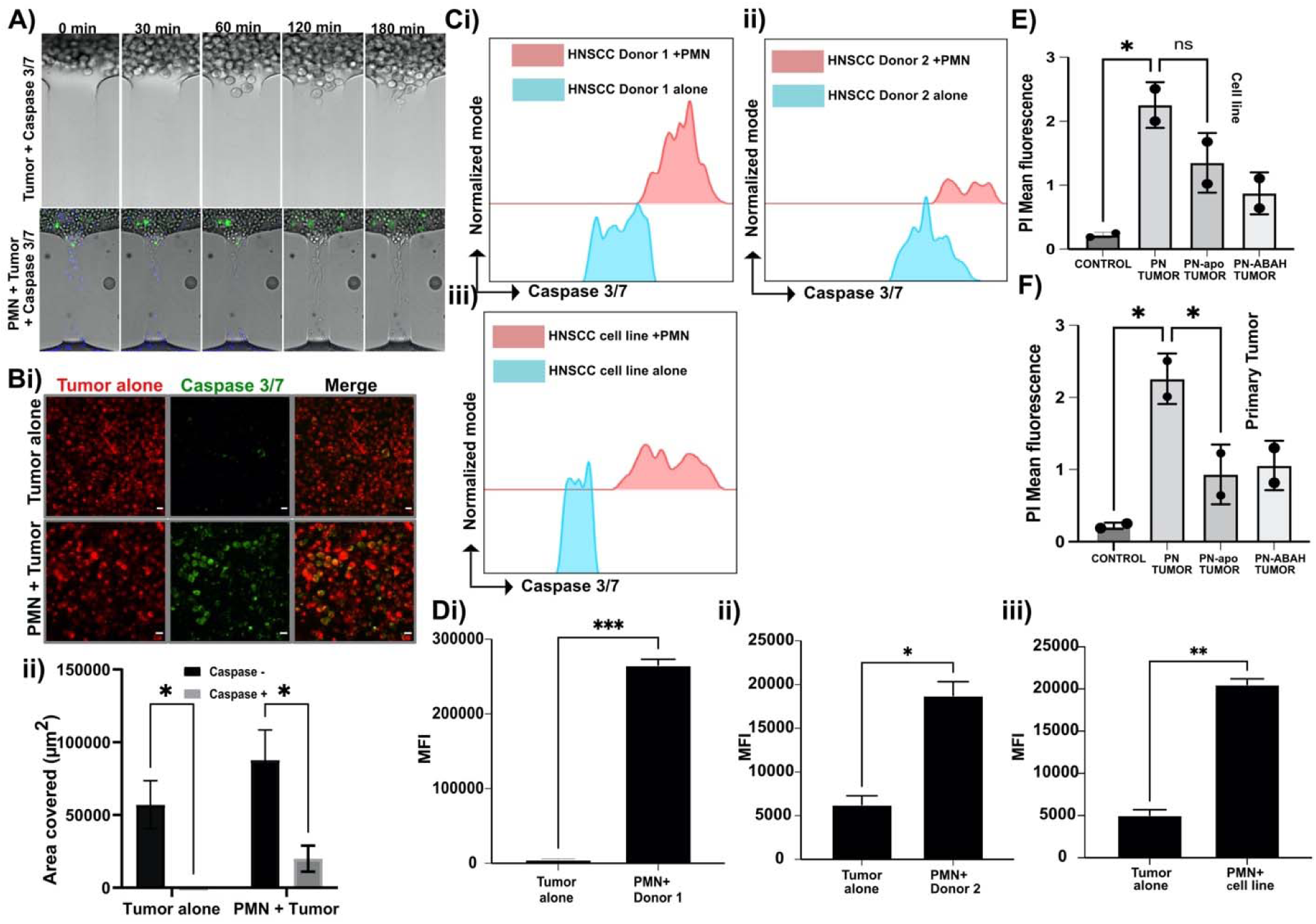
Neutrophil trigger caspase 3/7 pathway in tumor cell. **(A)** Sequential time microscopy images showing the appearance of caspase 3/7 event dye (green) in MDA-231 tumor cells (non GFP) in response to primary neutrophils cluster (blue) in tumor alone and tumor + neutrophils at 0 min, 60 min, 120 min, 240 min. **(Bi)** Microscopy image of caspase 3/7 event dye (green) positive primary tumor cells in the presence and absence of neutrophils. Images are representative of 5 independent experiments. To confirm tumor death, we quantified **(Bii)** the area covered by caspase positive tumor cells in tumor alone and neutrophil tumor coculture. *P* values were determined by two-way ANOVA; \**P*□<□0.05. **(C)** Representative flow cytometry histograms of caspase 3/7 staining on tumor cells after 24 h incubation with neutrophils using **(i-ii)** two different donors and **(iii)** tumor cell line. (Di-iii) Median fluorescence intensity (MFI) of flow cytometry analysis in **C (n=3).** *P* values were determined by Unpaired t test (parametric test); \**P*□<□0.05 and \*\**P*□<□0.01. Using fluorescent microscopy, apocynin (apo) and MPO inhibitor (ABAH) reduces the cytotoxicity of neutrophil against **(E)** primary tumor and **(F)** tumor cell line. Comparison between the area covered by caspase positive and negative tumor cells were performed by two-way ANOVA; \**P*□<□0.05.

Next, to further evaluate and confirm neutrophil cytotoxicity more directly, we used flow cytometry analyses. Neutrophils are terminally differentiated leukocytes derived from the myeloid lineage. Therefore, we isolated neutrophils from peripheral blood using negative selection and confirmed their identity using the myeloid cell marker, CD11b (Supplementary Figure 3A). Tumor cells and isolated neutrophils were stained with vibrant red dye and cell tracker blue respectively prior to coculture. After 24 hours of neutrophil tumor coculture, cells were gently detached using trypsin, washed and resuspended in FACS buffer containing DNase 1 and flow cytometry was performed. Flow cytometry analysis was performed using the gating strategy (*See ‘gating strategy’ under material and methods*) (Supplementary Figure 3B). Applying this gating strategy, we identified cells (Supplementary Figure 3Bi) and single cells by excluding doublets (Supplementary Figure 3Bii) and then identified the tumor cell population (vibrant red dye) by gating on the cell population positive for vibrant red dye and excluding neutrophils (cell tracker blue) (Supplementary Figure 3Biii). Next, we quantified the number of vibrant red dye positive cells that were undergoing cell death based on their expression of apoptosis marker Caspase 3/7 (Supplementary Figure 3Biv & v and C).

Both microscopy imaging and flow cytometry analysis revealed that in the presence of neutrophils, patient-derived tumor cells (Figure 5B, Ci-ii and Di-ii) and tumor cell lines (Figure 5Ciii and 5Diii) underwent increased cell death based on their expression of apoptosis marker Caspase 3/7 compared to tumor alone. To probe into the molecular mechanisms regulating neutrophil-mediated tumor cell death via microscopy, we used chemical inhibitors to target the ROS production pathway including apocynin, known to inhibit NADPH oxidase and 4-aminobenzoic acid hydrazide (4-ABAH) which inhibits myeloperoxidase (MPO). We observed that in the presence of Apocynin and 4-ABAH, there were less PI-positive tumor cell line (Figure 5E) and primary tumor cells (Figure 5F) compared to the untreated neutrophil group.

### Tumor spheroids recruit neutrophils and NK-92 cells *in vitro*

The TME is characterized by a network of cancer cells and recruited immune cells. However, the interaction among the different immune cells is still not well understood. Natural killer (NK) cells are innate lymphoid cells that play critical roles in protecting against tumor invasion and development (*36*). In human patients, a higher number of circulating or tumor-infiltrating NK cells is correlated with better patient outcomes (*37*). To better understand the immune cell-immune cell interactions during tumor progression, we sought to study the interaction between neutrophils and NK cells in terms of migration using a tumor spheroid to mimic tumor mass in culture.

To do this, we first generated multicellular tumor spheroids using patient derived tumor cells with diameters ranging from 100 to 200 µm using the hanging drop method (*17*). Next, we investigated the recruitment ability of the multicellular tumor spheroid by co-culturing the tumor spheroid with neutrophils isolated from healthy donors and/or NK-92 cells. We used NK-92 cells because they are a human NK cell line and are known for retaining cytotoxic capacity both *in vitro* and *in vivo* and have been extensively tested in clinical trials (*38*). We co-cultured multicellular tumor spheroids with NK-92 cells (1 spheroid:10^4^ NK-92 cells) or neutrophils (1 spheroid: 10^5^ cells) for 24 h. Microscopy analysis showed that both immune cell types migrated toward the multicellular tumor spheroid (Figure 6A-D).

**FIGURE 6:**
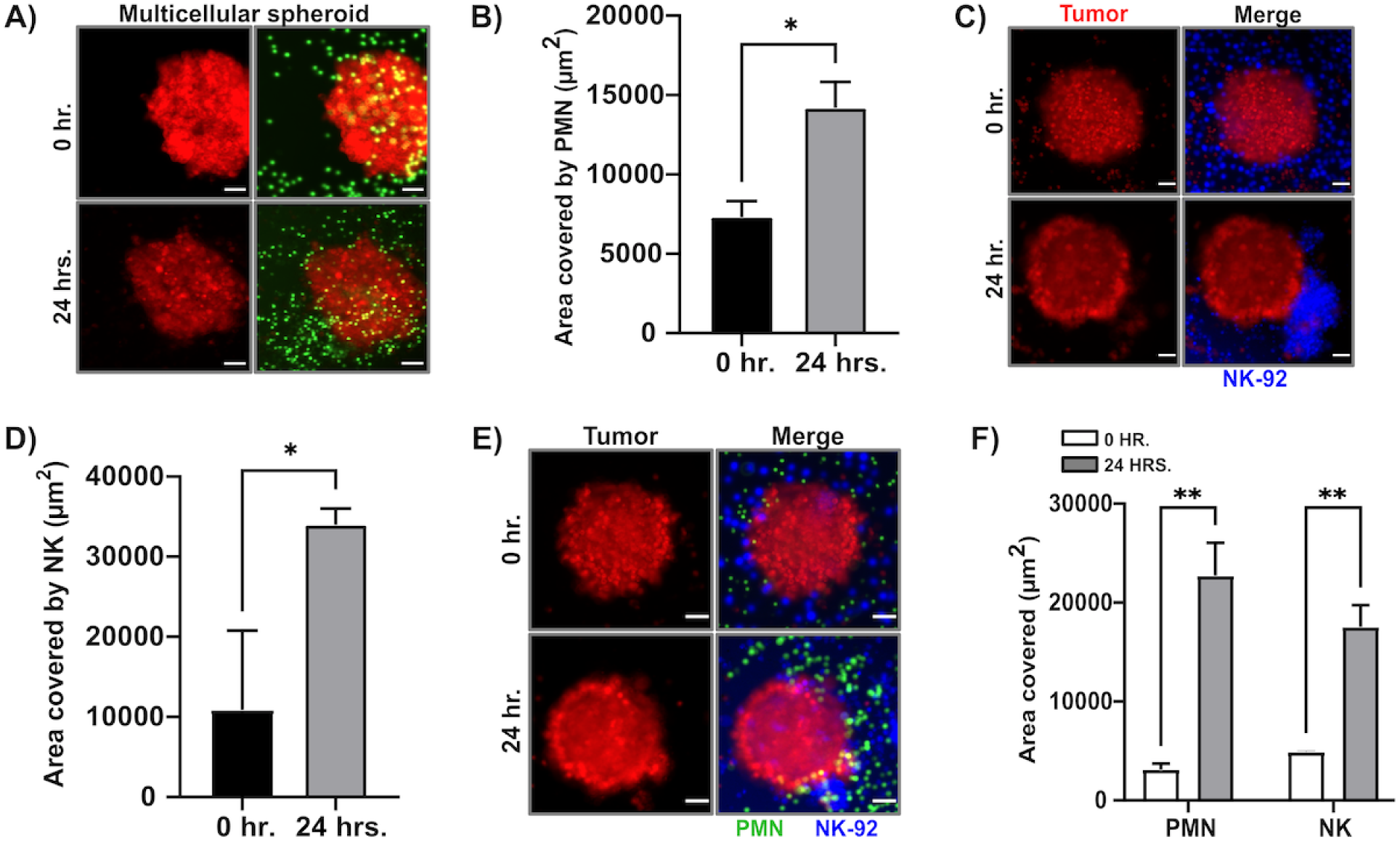
Tumor spheroids recruit neutrophils *in vitro*. Representative microscopy image showing clusters of primary neutrophils around **(A)** multicellular tumor cell spheroid. **(B)** Quantification of area covered by neutrophils around the multicellular tumor cell spheroid. Representative microscopy image showing clusters of NK-92 cells around **(C)** multicellular spheroid. **(D)** Quantification of area covered by NK-92 cells around the multicellular tumor cell spheroid. Representative microscopy image showing clusters of neutrophils and NK/92 cells around **(E)** multicellular spheroid. **(F)** Quantification of area covered by NK-92 and neutrophils. Each bar represents mean ± standard deviation from N= 3 (primary tumor cell spheroid) or 3 (Multicellular spheroid) independent donors. Comparison between the area covered by NK and PMN in 0 hr. and 24 hrs. were performed by unpaired t test; \**P*□<□0.05 (B and D) and comparison between the area covered by NK and PMN in 0 hr. and 24 hrs. were performed by two-way ANOVA; \*\**P*□<□0.01.

To investigate the interaction between the different immune cell types during tumor progression, we performed a triple coculture using primary neutrophils, NK-92 cells, and multicellular tumor spheroids for 24 h. Primary neutrophils, NK-92 cells, and multicellular tumor spheroids were stained with calcein, cell tracker blue and vibrant red dye respectively to confirm migration. Thus, we co-cultured primary neutrophils with multicellular tumor spheroids for 2 hours to mimic neutrophils arriving earlier at the tumor site which would be followed by NK-92 cells. Both neutrophils and NK-92 cells migrated toward the multicellular tumor spheroid in our triple coculture assay (Figure 6E). Microscopy analysis revealed clusters of neutrophil around the multicellular tumor spheroid and the area covered by neutrophils and NK-92 cells was ∼□5 and 4 times larger after 24 h. respectively (∼□500 μm^2^ vs. ∼□25,000 μm^2^, at 0 h. vs. 24 h. for neutrophils; ∼□500 μm^2^ vs. ∼□20,000 μm^2^, for NK-92 cells) (Figure 6F).

### Tumor cells induce changes in neutrophil gene expression associated with killing and migratory response *in vitro*

Next, we sought to evaluate the molecular changes in neutrophils in response to interacting with the tumor cells and NK cells. In this context, we investigated the activity of neutrophil-related genes involved in killing in our *in vitro* assay. Neutrophils contain different types of granules. The primary granules contain histotoxic enzymes, including elastase (NE), MPO, and antimicrobial enzymes, cathepsins, defensins and lactoferrin (*39*). Neutrophils were cocultured with tumor cells and NK cells or alone (control). After 24 h. in coculture, we used a negative selection strategy to deplete NK and tumor cells, using anti-CD56 conjugated magnetic beads to deplete NK-92 cells and anti-EpCAM conjugated magnetic beads to deplete tumor cells. Depletion of NK and tumor cells left a purified population of neutrophils in suspension. We then performed reverse transcription quantitative polymerase chain reaction (RT-qPCR) on the isolated neutrophils. RT-qPCR experiments and nonhierarchical clustergram analysis showed that neutrophil-related genes were up regulated in the iPMN compared to neutrophils alone (Figure 7A). More specifically, we observed that genes associated with neutrophil killing (MPO, NOX-1, LTF and NE) were all significantly up-regulated in iPMN compared to neutrophils alone (Figure 7A-B).

**FIGURE 7:**
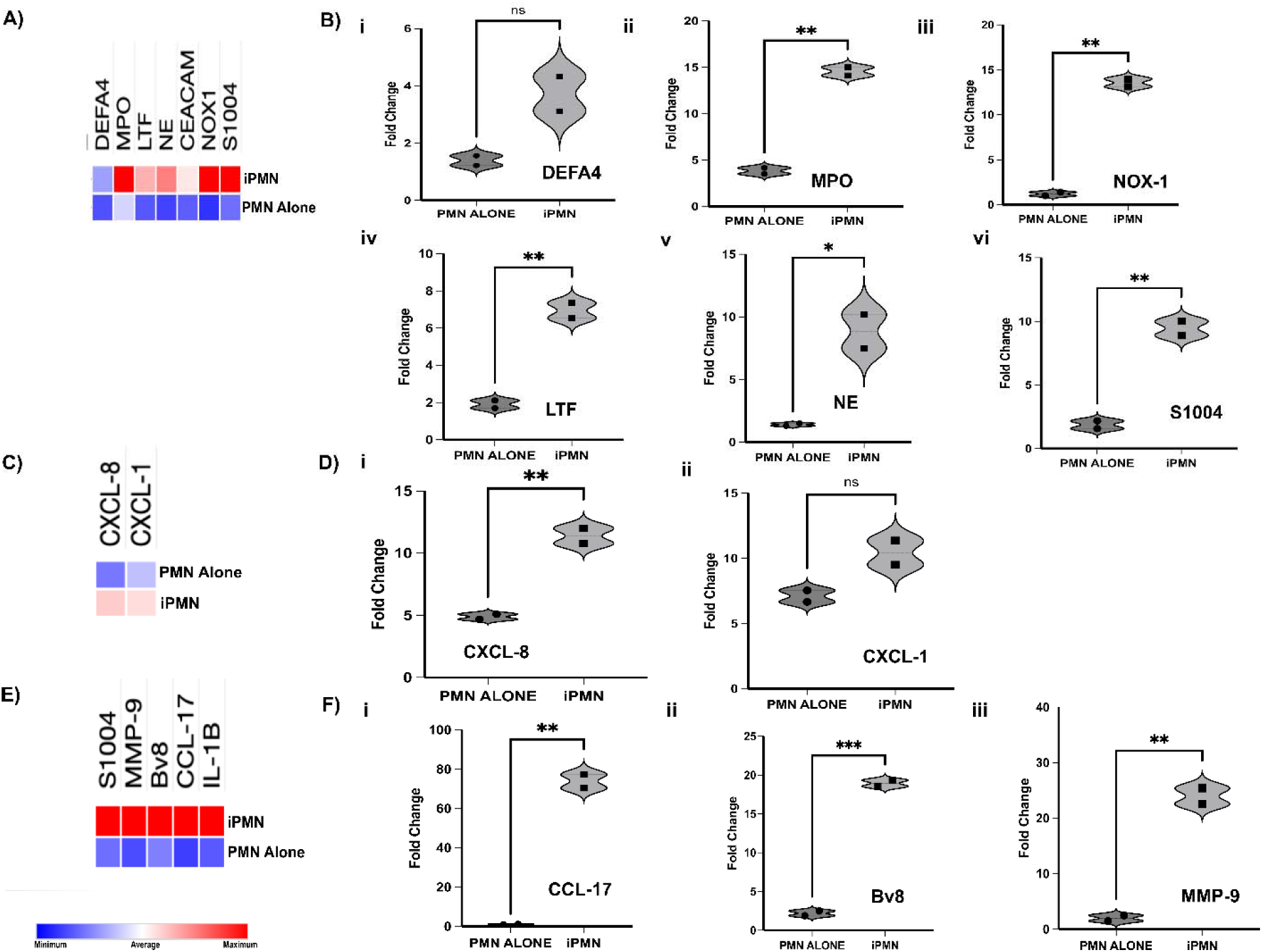
Tumor cells induce killing and migratory responses in neutrophils *in vitro.* Primary neutrophils were isolated from the co culture (iPMN) and RT-PCR was carried out to analyze their gene expression. **(A)** Cluster gram comparing gene expression associated with killing in neutrophils in iPMN with neutrophils alone and LPS stimulated neutrophils. **(B)** Bar graphs showing upregulation of multiple genes associated with killing in neutrophils in iPMN compared with PMN alone. **(C)** Cluster-gram comparing gene expression associated with migration in neutrophils in iPMN with neutrophils alone. **(D)** Bar graphs showing upregulation of genes associated with migration in neutrophils in iPMN compared with PMN alone. **(E)** Clustergram comparing gene expression associated with immunosuppression in neutrophils in iPMN with neutrophils alone. **(F)** Bar graphs showing upregulation of genes associated with immunosuppression in neutrophils in iPMN compared with PMN alone. Graphs show means ± SD. Three biologically individual experiments were carried out, but data represented as mean ± SD of 2 independent donors. Comparison of gene expression between PMN and iPMN were performed by unpaired t test; \**P*□<□0.05, \*\**P*□<□0.01 and \*\*\**P*□<□0.001.

Numerous studies have suggested that soluble factors, such as CXCL chemokine family members, promote neutrophil infiltration in a variety of tumors (*40*–*42*). Our data also confirmed that genes associated with neutrophil recruitment such as CXCL-8, which encodes for a major IL-8 protein, accompanied by the induction of CXCL-1 expression (∼10-fold increase) was upregulated in iPMN compared to neutrophils alone, suggesting that neutrophils are recruited toward tumor cells (Figure 7C-D). On the other hand, genes associated with tumor invasion and metastasis (MMP-9 and CCL-17) were also significantly up-regulated in iPMN compared to neutrophils alone (Figure 7E-F), suggesting a possible pro-tumor role of neutrophil in the TME. Together, these results indicate that tumor cells can induce a pro-inflammatory response in neutrophils as well as pro-tumor response *in vitro*.

### Neutrophils suppress NK tumor killing *in vitro*

Previous studies have demonstrated the immuno-suppression of T cells which can be driven by several neutrophil derived immunoregulatory factors like NO and ROS **(*43*)**. However, the effect of the interaction between neutrophils and NK cells on tumor killing remains unclear. Thus, we sought to investigate the effect of neutrophils on NK-92 tumoricidal activities *in vitro*. To do this, we first investigated if NK-92 cells alone could exert tumor death *in vitro* by coculturing NK-92 cells and patient derived tumors at the ratio of 1 tumor cell: 2 NK cells for 24 h, followed by the addition of caspase 3/7 and/or PI. We observed that NK-92 exerted more primary tumor cells (Figure 8A) and tumor cell line (Figure 8B) death after 24 h by measuring the PI fluorescence via microscopy compared to neutrophils, thus providing a baseline for our killing assay *in vitro*.

**FIGURE 8:**
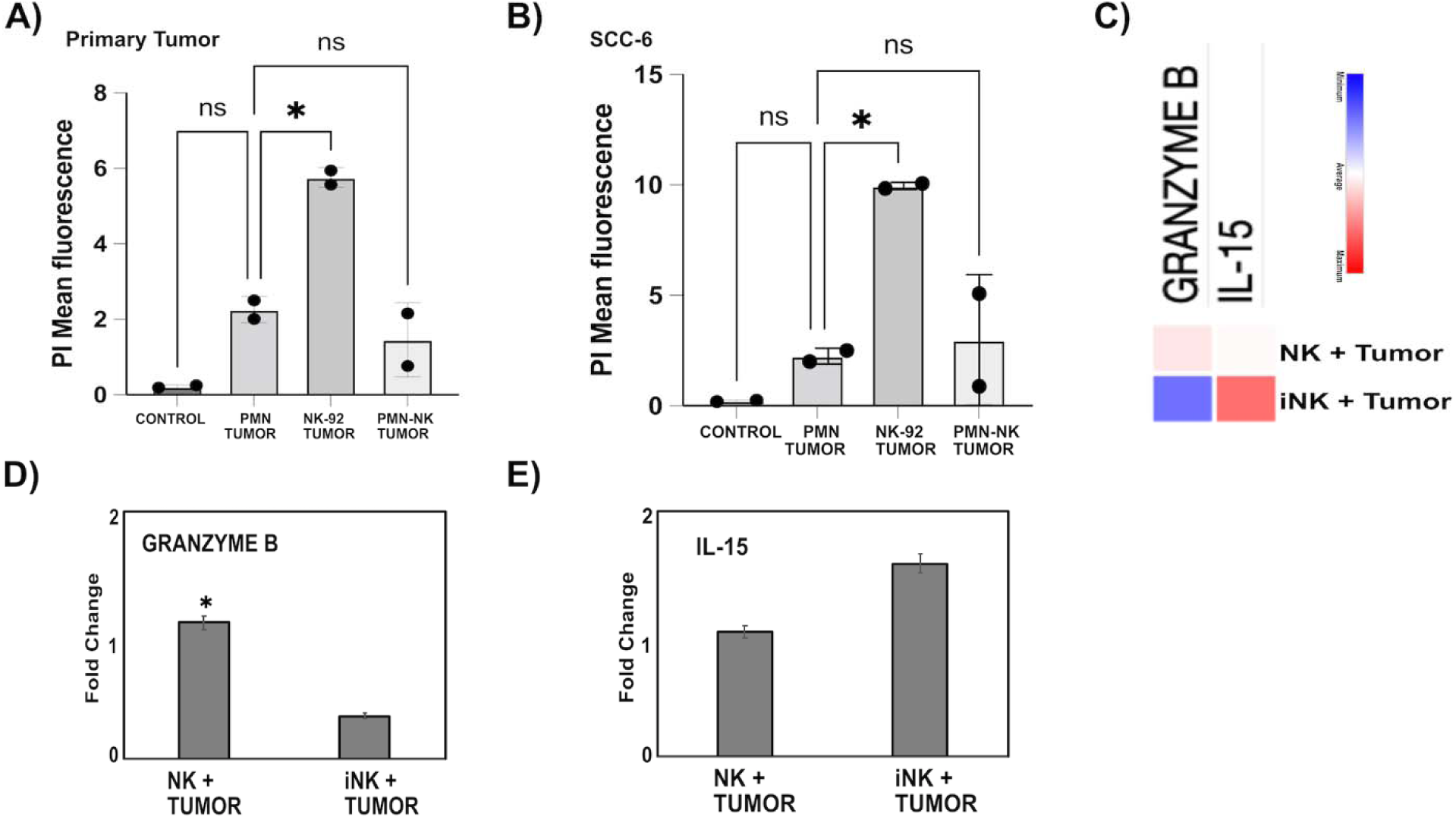
Neutrophils suppress NK-92 tumoricidal effect *in vitro*. Graphs show the mean PI fluorescence of **(A)** primary tumor cells and **(B)** tumor cell line in our well plate culture. **(C)** Cluster gram comparing gene expression associated with killing and survival in isolated NK-92 from triculture (iNK-92) with NK-92 alone. Bar graphs showing downregulation/upregulation of genes associated with **(D)** killing and **(E)** survival in iNK-92. Comparison between the area covered by PI positive and negative tumor cells were performed by two-way ANOVA; \**P*□<□0.05. Comparison of gene expression between NK and iNK were performed by unpaired t test; \**P*□<□0.05.

To examine if neutrophils could increase NK-92 cell cytotoxicity *in vitro*, we carried out a triple coculture of neutrophils, NK-92 cells and patient-derived tumor cells. To do this, we first cocultured a confluent monolayer of patient-derived cancer cells with primary neutrophils for at least 1 h. to mimic neutrophils arriving earlier at the tumor site followed later by NK-92 cells. Microscopy analysis of caspase 3/7 fluorescent area revealed that co-culture of neutrophils with NK-92 cells did not increase NK-92 tumoricidal effects, but rather it reduced the tumoricidal ability of NK-92 cells, which could suggest the involvement of neutrophil-mediated NK-92 cell cytotoxicity suppression *in vitro* (Figure 8A-B).

To further quantitatively confirm our observation, we employed flow cytometry (using caspase 3/7) (Supplementary Figure 4). Natural killer cells are members of the innate lymphoid cell (ILC) family and characterized in humans by expression of the phenotypic marker CD56^+^ and CD3^-^. Therefore, we used CD56 to identify NK-92 cells (Supplementary Figure 3A). In a parallel experiment, tumor cells and NK-92 were stained with vibrant red dye and cell tracker CMTPX respectively prior to coculture. After 24 h coculture, cells were detached using trypsin, washed and resuspended in FACS buffer containing DNase 1 and flow cytometry was performed. Flow cytometry analysis was performed using similar gating strategy shown and described in Supplementary Figure 4A. Flow cytometry analysis revealed that in the presence of NK-92 cells, patient-derived tumor (Supplementary Figure 4Bi-ii & 4Ci-ii) underwent cell death based on their expression of apoptosis marker Caspase 3/7 compared to the neutrophil group.

Next, we wanted to examine the molecular changes in NK-92 driven by neutrophil-released ROS *in vitro*. We isolated NK-92 cells from our triple coculture (iNK-92) to investigate the expression of NK-92 cell-related genes crucial to NK cell killing and survival. Briefly, we isolated NK-92 from the tri-culture using anti-CD56 magnetic beads. We then carried out a RT-qPCR analysis to analyze the expression of Granzyme B (GZMB) and IL-15 on the isolated NK-92 cells. NK cells eliminate virus-infected and tumor cells by releasing cytotoxic granules containing GZMB. On the other hand, IL-15 promotes NK cell survival via maintenance of the antiapoptotic factor Bcl-2. RT-qPCR analysis revealed that GZMB was significantly downregulated and IL-15 was slightly up regulated in NK-92 from the tri-culture (iNK-92) compared to NK-92 alone supporting the suppression in the killing ability in NK-92 cells generated by the presence of neutrophils (Figure 8C-E). The gene expression analysis further demonstrated that neutrophils were indeed suppressing cytotoxicity in NK-92 *in vitro*.

### NETs shielded tumor spheroid are protected from NK-92 cytotoxicity and migration

Having shown the presence of NETs in our co-culture and breast cancer human tissue, we sought to investigate the possible functional effect of the released NETs on the surrounding immune cells in the TME. To investigate the possible immunomodulatory effect of NETs in NK cell responses, we generated a ring of NETs around a tumor spheroid *in vitro* (Figure 9A). Tumor spheroids were generated from patient derived tumor cells using the hanging drop method. Next, we co-cultured the tumor spheroids with healthy donor neutrophils in the presence and absence of phorbol 12 myristate 13-acetate (PMA) to induce NET formation. In the absence of NETs, we observed that NK-92 cells migrated and penetrated into the tumor spheroid (Video 3). In the presence of PMA, NETs were released and were stained with sytox green (impermeant nucleic acid fluorescent dye) and confirmed via confocal microscopy (Figure 9B-C and Video 4). To examine whether we could enhance NK-92 cell migration and contact with the tumor spheroid, we added DNase 1 in the assay (DNase 1 is aimed at cleaving the DNA fragments) (Figure 9D-E). Inclusion of DNase reduced the observed sytox green staining to nearly negligible levels, demonstrating it was indeed degrading the NETs (Figure 9D-E). Using confocal microscopy, we observed that the number of NK-92 cell penetration into the tumor spheroid was reduced in the presence of NETs (Figure 9F). However, the treatment with DNase I trended towards increased number of penetrating NK-92 cells (Figure 9F & 9G). Microscopy analysis of NK-92 recruitment by fluorescence intensity showed that in the presence of NETs, NK-92 cells demonstrated oscillatory migratory behavior compared to without NETs (Figure 9H). Taken together, our data suggest that NET formation could also be contributing to tumor survival by protecting tumor cells from NK cytotoxicity.

**FIGURE 9:**
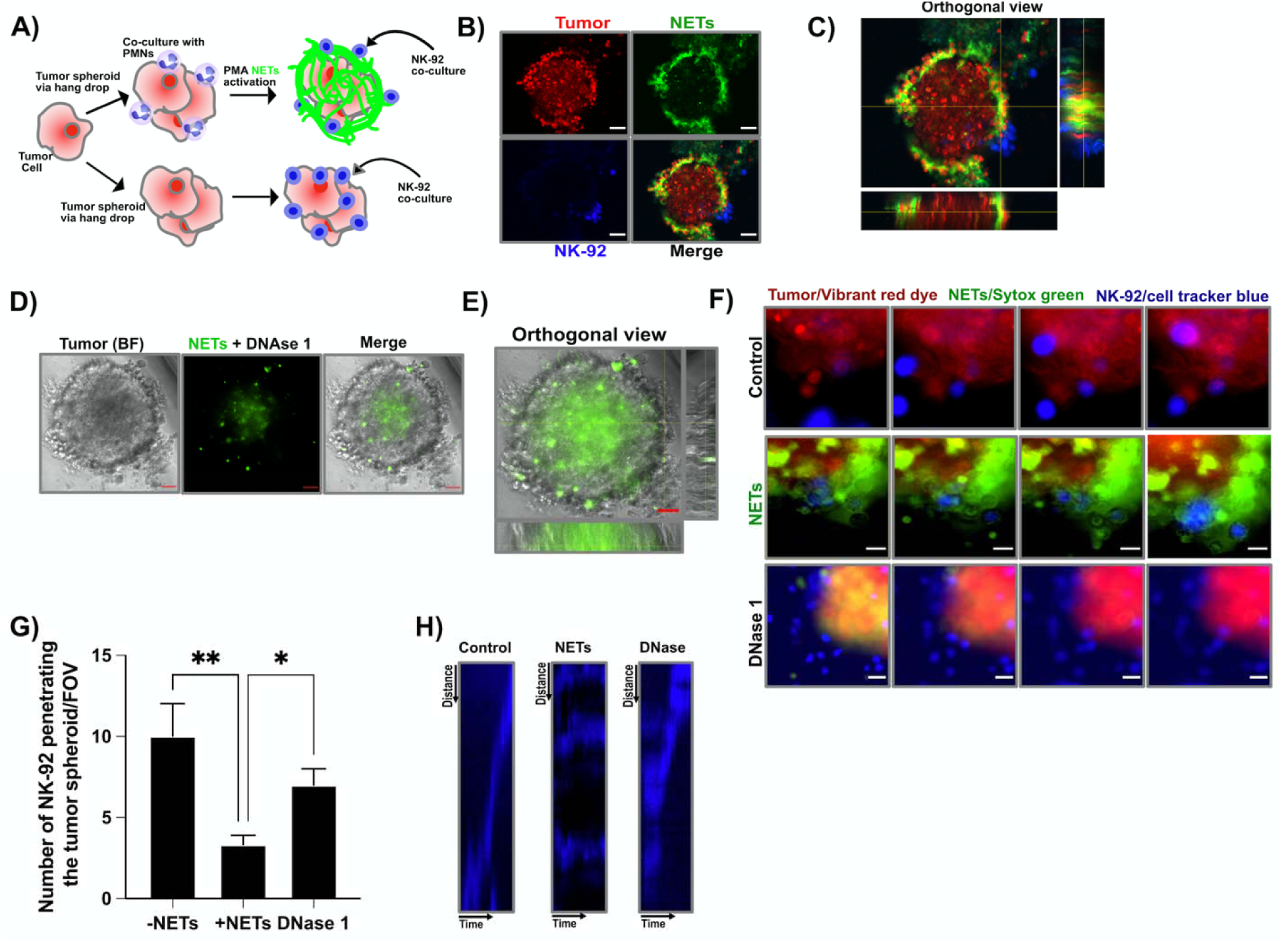
NETs inhibit NK-92 cytotoxicity and migration *in vitro.* **(A)** Schematic illustration of experiments in **(D)** to **(H).** Tumor spheroids were labeled with vibrant red dye (red) and cultured in the presence of neutrophils plus **(B and C)** PMA (NET) and **(D and E)** DNase I (DNase). Tumor spheroids were subsequently co-cultured with NK-92 cells. **(F)** Representative frames of videos of cytotoxic cells (blue) co-cultured with tumor spheroids (red) in the absence or presence of NETs (green) as schematized in **(A).** (**G**) Graph showing the number of NK-92 cells penetrating tumor spheroid in the presence and absence of NETs per field of view. P values were determined by ordinary one-way ANOVA; \**P*□<□0.05 and \*\**P*□<□0.01. **(H)** Kymograph of migrating NK-92 in the presence and absence of NET.

## DISCUSSION

The TME is a complex system in which stromal cells and several types of immune cells often play a dual role reducing or promoting tumor growth, invasion, and metastasis (*44*). Several immune cells such as macrophages, T cells, and NK cells, can directly or indirectly influence tumor growth and progression by secreting factors and molecules such as vascular endothelial growth factor (VEGF), transforming growth factor (TGF-beta) and matrix metalloproteinases (MMPs) that exert protumor or antitumor functions (*45*). The effect of neutrophil-tumor interaction on tumor progression still remains unclear. Here we have demonstrated that naive neutrophils interact with tumor cells and that they can exert restriction on tumor metastasis *in vitro*. These findings validate prior published data (*46*, *47*) but also offer a substantial expansion of knowledge by demonstrating that neutrophils from healthy donors can restrict tumor metastasis through their swarming response and their effector functions (NETs and ROS). On the other hand, our findings also demonstrate that neutrophil effector functions can concurrently inhibit NK cell cytotoxicity via down regulation of granzyme B expression.

In the current study, we used a simple *in vitro* assay and a UOM-based *ex vivo* assay to investigate the interactions between naive primary neutrophils and tumor cells. We observed that neutrophils form clusters when added on top of a tumor monolayer or cultured with tumor spheroids. Those observations led us to explore the effect of their interaction on tumor killing using patient derived HNSCC tumor epithelial cells and an HNSCC cell line as target cells. We found that the interaction between neutrophils and the tumor cells has the tendency to be both protumor and antitumor as it can both induce tumor death through ROS production and promote tumor progression via NETs release. The optical metabolic imaging data reveals alterations in the metabolic activity of neutrophils during the various phases of swarming in response to tumor cells. Importantly, these observed changes in the metabolic activity are particularly pronounced in neutrophils that are close to tumor cell(s). Our data further show that primary neutrophils had a higher initial tumoricidal activity than NK-92 cells within 24 h, however, in our culture model, most of the neutrophils were dead after 24 h. We predict that NK-92 cells can sustain their cytotoxicity for longer period than primary neutrophils as they undergo suicidal NETosis in order to kill. On the other hand, our data also show that neutrophil effector functions may reduce NK-92 cell cytotoxicity, highlighting their pro-tumor and anti-tumor effects. In agreement with our findings, high concentrations of ROS are toxic to both tumor cells and immune cells (*48*).

Neutrophils may influence tumor progression through the paracrine release of cytokines and chemokines with protumor or antitumor functions, depending on the tumor microenvironment (*49*). Thus, we proposed that there might be cytokines or chemokines induced by the interaction between neutrophils and tumor cells that may in part account for the observed reduced NK cell cytotoxicity. In order to test that hypothesis, we performed RT-qPCR array experiments to investigate gene expression and found the expression of several pro-inflammatory genes such as CXCL8 and IL1B, killing related genes such as DEFA4, NOX-1, MPO, MMP-9 and NE and neutrophil suppression genes like Bv8, MMP-9 and CCL-17 were all upregulated after neutrophil-tumor co-culture.

CXCL8 chemokine is a major regulator of neutrophil migration (*50*), which may mediate neutrophil infiltration into TME. Our study shows that CXCL-8 expression was increased after neutrophil-tumor co-culture. In agreement with our finding, recruited neutrophils in the TME have been shown to express CXCL1 and CXCL8, thereby inducing tumor angiogenesis and subsequent tumor progression and metastasis (*51*). In agreement with our finding, CCL17, is a known neutrophil derived chemokine that binds to CCR4 and expressed in T-helper type 2 (Th2) cells and in regulatory T cells (Tregs) (*52*). Consistently, TANs have been shown to recruit regulatory T cells (Tregs) in a mouse model of cancer, mainly *vi*a CCL17 (*53*). Several lines of evidence indicate that infiltrating neutrophils can promote metastasis by producing growth factors such as Bv8 (*54*). We confirmed that Bv8 was abundantly expressed in iPMN compared to neutrophil alone and LPS activated neutrophils, which further support Bv8 expressing neutrophils may contribute to tumor progression.

Neutrophils contain different types of granules. The primary granules contain histotoxic enzymes, including elastase (NE), MPO, and antimicrobial enzymes, cathepsins, defensins and lactoferrin (*55*). Molecular analysis demonstrated that genes associated with killing in neutrophils such as, NOX-1, MPO, and NE were up-regulated in iPMN from tumor cocultures compared to neutrophils alone. Interestingly, PCR analysis revealed changes in pathways associated with killing in iNK-92. Our molecular analysis demonstrated that granzyme B was down-regulated in iNK-92 compared to NK-92 alone. In this context, our data revealed that the presence of neutrophils led to reduced activation and cytotoxicity ability in NK-92 cells. However, inhibition of ROS in our triple co-culture, partially restores NK-92 cytotoxicity, suggesting that this mechanism is involved in neutrophil mediated NK-92 suppression. These insights into neutrophil-NK interactions within the tumor microenvironment lay important groundwork for improving our understanding of the complex interplay between immune cells in this environment.

NETs are composed of DNA coated with histone, NE, MPO and MMP-9 (*56*). NETs can further trap circulating tumor cells (CTCs) *in vitro*, thereby facilitating their migration into distant organs (*57*). NETs have been observed in clinical samples of triple-negative breast cancer patients, driving tumor metastasis. To further investigate the role of NETs in TME, we encapsulated tumor spheroid with NETs to check if it reduces NK-92 migration *in vitro*. Our data suggests that NETs may be responsible for shielding tumor spheroids against NK-92 mediated cytotoxicity. However, the NETs mediated protective mechanism was reduced when the coated NETs were digested using DNase. Interestingly, the action of DNase appears to have significant effect on migration and contact of NK-92 cells with the tumor spheroid. However, it should be noted that our work does not necessarily imply that the physical barrier created by NETs is the sole mechanism linked with the reduction of NK-mediated immunity. There may be alternative mechanisms that are currently unknown. In support of our finding, NETs have been implicated and observed to be surrounding a metastatic tumor model in the lung and in the liver sinusoids (*58*). In fact, DNase treatment has been shown to efficiently degrade NETs and attenuate the development and progression of liver metastases in a murine model of colorectal cancer (*59*). Clinically, NETs have been observed in clinical samples of triple-negative breast cancer patients as well as in *in vivo* mouse models even in the absence of infection. NET digestion with DNase I markedly reduced lung metastases in mice, indicating the crucial roles of NET in metastasis even in the absence of infection (*60*). Taken together, we further confirm the double role of neutrophils at tumor site and extend these findings into human cells.

Previous reports have studied the suppression of NK cytotoxicity mediated by neutrophils *in vivo* (*61*). In this study, we specifically and quantitatively confirmed tumor cell death and the suppressed tumoricidal ability in NK-92 driven by neutrophils using both microscopy and flow cytometry techniques. We further analyze the molecular changes in neutrophils that may be responsible for the observed NK cell dysfunction. One limitation of this study is that we used RT-qPCR to analyze **∼**20 genes related to both neutrophil and NK cell response, which only represents a fraction of the total amount of alterations observed *in vivo*.

Future studies could leverage RNA-sequencing technologies to analyze thousands of genes to provide a more in-depth molecular analysis of the suppression process. Collectively, our work identifies an immune-associated cellular and molecular network involving neutrophils and NK cells which interact at the tumor site and may contribute to tumor progression. These findings support further investigation into neutrophils as potential therapeutic targets for cancer treatment that could ultimately improve patient outcomes.

## Supporting information

Tumor alone migration

PMN alone migration

PMN Tumor interaction

PMN alone + Sytox green dye

PMN + Tumor + Sytox green dye

Tumor alone + Caspase 3/7 dye

PMN + Tumor + Caspase 3/7 dye

NK-92 cells migrating toward tumor spheroid

NK-92 cells + NETs + Tumor spheroid

NK-92 cells + NETs + DNAse 1

## ACKNOWLEDGEMENTS

This work was supported by the University of Wisconsin Carbone Cancer Center, Cancer Center Support Grant NIH P30CA014520. This work was also supported by the National Institutes of Health NIH R01AI134749 to AH and DB, the National Institutes of Health NCI R01 CA085862 to AH, the National Institutes of Health NIH U24AI152177 to DB and SK, and the Swiss National funding (P500PB_203002) and Jubilaumsstiftung von Swiss Life (project number: 1350) to K.A.B. We thank Alice Golubiewski in the Beebe Lab for arranging whole blood collection from healthy donors. We also thank Chao Li for the discussion and training on the fabrication and use of the under oil microfluidic assay. Finally, we thank the UW Optical Imaging Core for the support on the confocal imaging.

## AUTHOR CONTRIBUTIONS

K.A.B designed and performed experiments and did data analysis with assistance from S.C.K and D.J.B. K.A.B and N.W.H prepared the microfluidic devices and J.A.M assisted with the tumor spheroids and NK cells biology. K.A.B and R.D performed the OMI experiment and R.D generated and analyzed the OMI data with assistance from M.C.S. K.A.B., R.D., J.A.M., A.H., M.C.S., D.J.B., and S.C.K. wrote and reviewed the manuscript.

## DATA AND MATERIALS AVAILABILITY

All data needed to evaluate the conclusions in the paper are present in the paper and/or the Supplementary Materials.

