## Supplementary Figure 1 for "Naive primary neutrophils play a dual role in the tumor microenvironment"

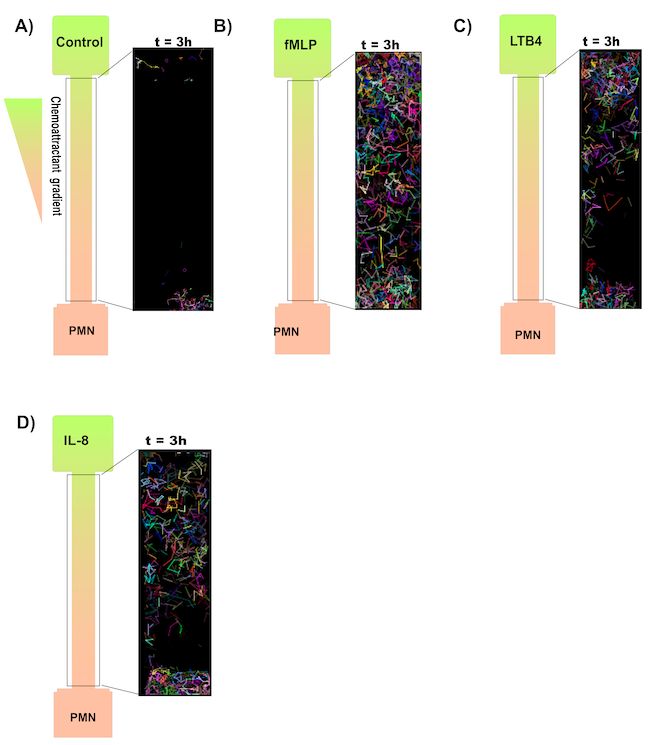

**Fig. S1.** Neutrophil migration in the UOM device. Schematic illustration showing migratory tracks of neutrophils migrating in A) Control B) *f*MLP gradient, C) LTB_4_ gradient and D) IL-8 gradient. All images were designed by Affinity Designer Software (version 1.10.6.) <https://affinity.serif.com/en-us/designer/>.
