## Supplementary Figure 2 for "Naive primary neutrophils play a dual role in the tumor microenvironment"

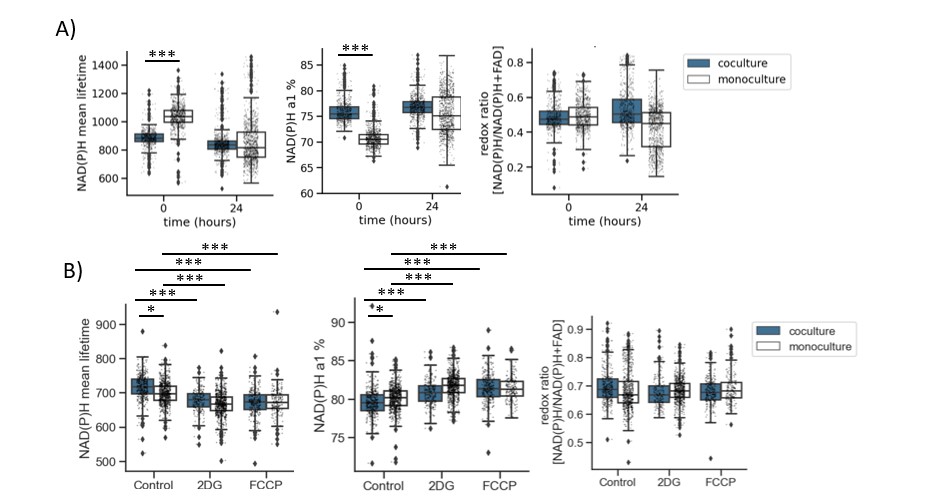

**Supplementary Figure 2:** Independent biological repeats of autofluorescence imaging of neutrophil tumor interaction. **A)** Quantification of NAD(P)H τ_m_, NAD(P)H α1% and optical redox ratios of neutrophils cultured either in monoculture or coculture with tumor cells before (0-1 hour) ‘0’ and after (24-25 hour) ‘24’ swarming response. Optical imaging data is analyzed for atleast n = 9 images per condition from the 2-3 independent dishes. The data are computed at the single-cell level and the number of cells segmented per condition is presented in Table S2. **B)** Quantification of NAD(P)H τ_m_, NAD(P)H α1% and optical redox ratios of neutrophils cultured either in monoculture or coculture with tumor cells either treated with 100mM 2DG, 660nM FCCP for 0-1 hours or no treatment (Control). Optical imaging data is analyzed for atleast n = 9 images per condition from the 2-3 independent dishes. The data are computed at the single-cell level and the number of cells segmented per condition is presented in Table S4. Statistical significance was accessed using one-way ANOVA with Tukey’s post hoc test (***P < 0.001; *P <0.5). Data is presented as medians and error bars are the 95% confidence interval.
