## Supplementary Figure 3 for "Naive primary neutrophils play a dual role in the tumor microenvironment"

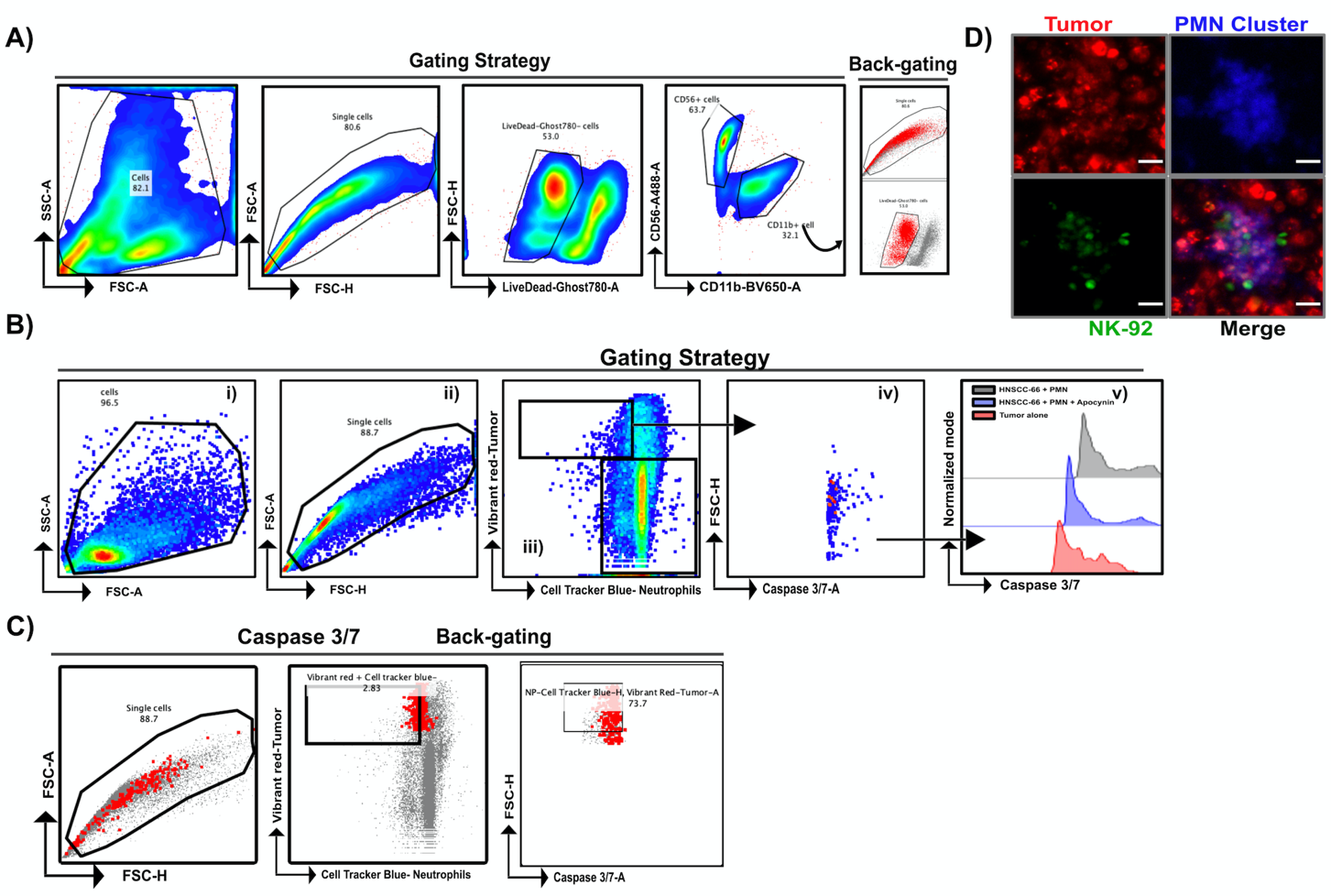

**Supplementary Figure 3:** Gating Strategies for identifying A) CD11b positive primary neutrophil, B) caspase 3/7 positive tumor cells and C) Back-gating strategies for caspase positive tumor cells D) Representative microscopy image showing primary neutrophil cluster (Hoechst dye-Blue) and NK cells (Calcein dye-Green) around tumor cells (vibrant red dye).
