## Supplementary Figure 4 for "Naive primary neutrophils play a dual role in the tumor microenvironment"

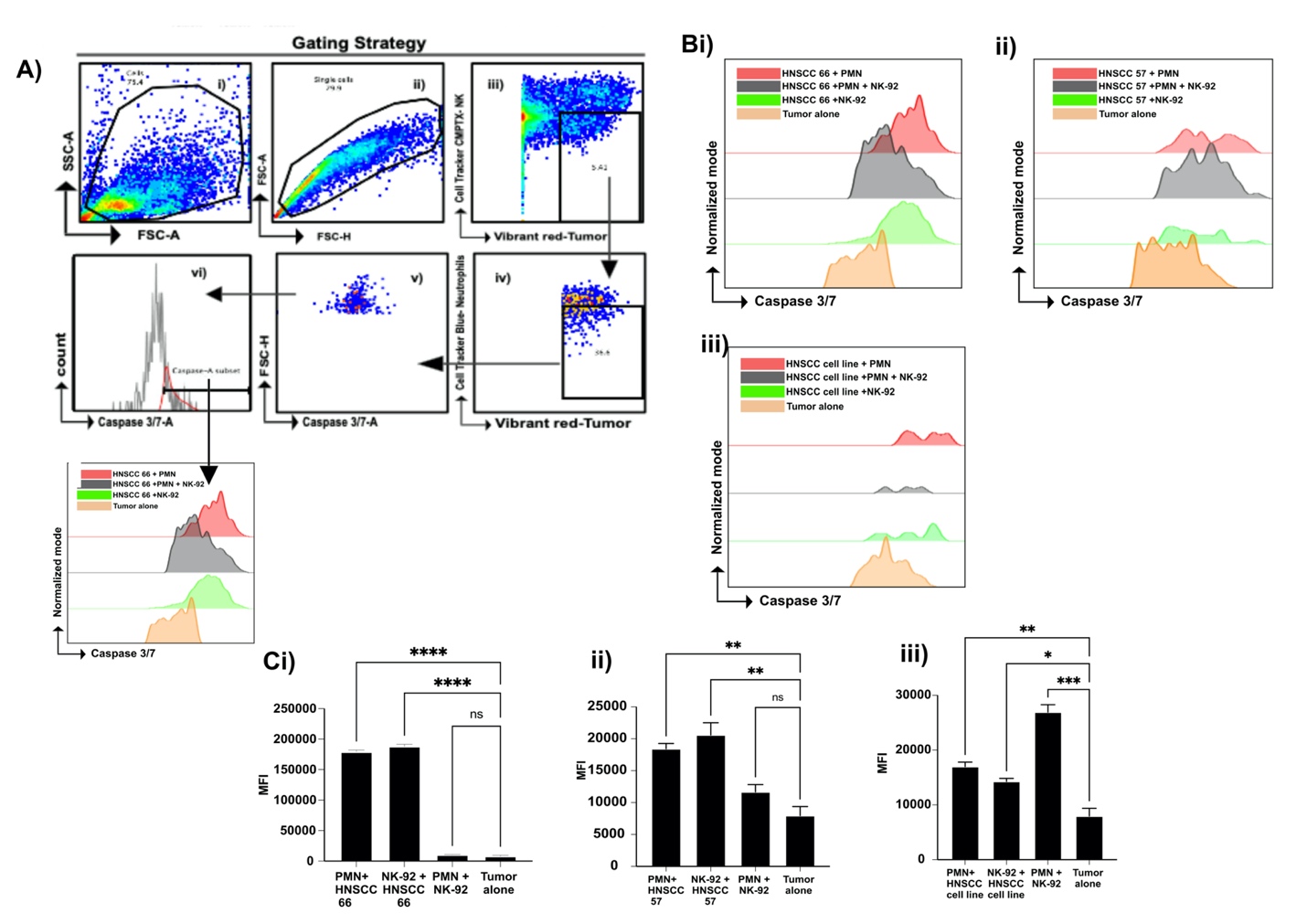
**Fig. S4.** Gating Strategies for identifying A) caspase 3/7 positive tumor cells. (B) Representative flow cytometry histograms of caspase 3/7 staining on tumor cells after 24 h incubation with neutrophils, neutrophil and NK-92 and NK-92 using **(i-ii)** two different donors **(iii)** tumor cell line. (C) Median fluorescence intensity (MFI) of flow cytometry analysis in **B (n=3).** *P* values were determined by ordinary one-way ANOVA; **P* < 0.05, ***P* < 0.01, ****P* < 0.001 and *****P* < 0.0001.
