## Supplementary Table 1 for "Naive primary neutrophils play a dual role in the tumor microenvironment"

**Supplementary Table 1:** Number of cells segmented for data shown in Figure (4B)

| **Time (hours)** | **Culture** | **Number of wells** | **Number of images** | **n = number of cells** |
| --- | --- | --- | --- | --- |
| 0 | Coculture | 3 | 9 | 512 |
|  | Monoculture | 3 | 9 | 868 |
| 6 | Coculture | 3 | 12 | 407 |
|  | Monoculture | 3 | 9 | 80 |
