## Supplementary Table 2 for "Naive primary neutrophils play a dual role in the tumor microenvironment"

**Supplementary Table 2:** Number of cells segmented for data shown in Supp Figure (3A)

| **Time (hours)** | **Culture** | **Number of wells** | **Number of images** | **n = number of cells** |
| --- | --- | --- | --- | --- |
| 0 | Coculture | 2 | 9 | 1386 |
|  | Monoculture | 3 | 9 | 820 |
| 24 | Coculture | 3 | 9 | 1768 |
|  | Monoculture | 2 | 9 | 1087 |
