## Supplementary Table 3 for "Naive primary neutrophils play a dual role in the tumor microenvironment"

**Supplementary Table 3:** Number of cells segmented for data shown in Figure (4D)

| **Time (hours)** | **Culture** | **Number of wells** | **Number of images** | **Group** | **n = number of cells** |
| --- | --- | --- | --- | --- | --- |
| 0 | Coculture | 3 | 9 | Contact | 68 |
|  |  |  |  | Non-contact | 444 |
| 6 | Coculture | 3 | 12 | Contact | 166 |
|  |  |  |  | Non-contact | 241 |
