## Supplementary Table 4 for "Naive primary neutrophils play a dual role in the tumor microenvironment"

**Supplementary Table 4:** Number of cells segmented for data shown in Supp Figure (3B)

| **Treatment** | **Culture** | **Number of wells** | **Number of images** | **n = number of cells** |
| --- | --- | --- | --- | --- |
| Control | Coculture | 3 | 9 | 377 |
|  | Monoculture | 3 | 9 | 698 |
| 2-DG | Coculture | 2 | 9 | 205 |
|  | Monoculture | 3 | 8 | 689 |
| FCCP | Coculture | 2 | 9 | 286 |
|  | Monoculture | 3 | 9 | 189 |
