## Supplementary Table 5 for "Naive primary neutrophils play a dual role in the tumor microenvironment"

**Supplementary Table 5:** Sequence of primers used for RT-PCR studies.

| **Gene** | Forward primer (5’-3’) | Reverse primer (5’-3’) |
| --- | --- | --- |
| EF-1 alpha | CAC CCA GGC ATA CTT GAA GG | GCC ACC TCA TCT ACA AAT GC |
| Defa4 | TGC AAG AGC AGA CCA TGC | GCC AGA AGA CCA GGA CAT |
| MPO | GGC ATC TCA CTG GAA CGG | CCT TTG ACA ACC TGC ACG AT |
| NOX-1 | AAA CAT TCA GCC CTA ACC AAA C | GAA TCT TCC CTG TTG CCT AGA |
| LTF | CCT GCC TCG TAT ATG AAA CCA C | AGC TGC ATA AAG AGA GAC TCC |
| ELANE | ATG TTT ATT GTG CCA GAT GCT G | GCA GGA CCC ACT GAG AAG |
| S100A4 | CAC GCC ATG ACA GCA GT | CTC TCT ACA ACC CTC TCT CCT C |
| CXCL-8 | CTT CAC ACA GAG CTG CAG AA | GAG ACA GCA GAG CAC ACA AG |
| CXCL-1 | GAA CAA GTC ATC CTC ATT GCC | CAG CCA ATC TTC ATT GCT CAA G |
| Bv8 | TCT TTC TTC TTT CCT GCC TTC C | GCT GTC AGT ATC TGG GTC AAG |
| MMP-9 | CGT CGA AAT GGG CGT CT | ACA TCG TCA TCC AGT TTG GTG |
| GZMB | CAG AGA CTT CTG ATC CCA GAT | TCC TGA GAA GAT GCA ACC AAT |
| IL-15 | TTT CCA GCA GCC ATC CAT C | GAG TTT GTT CCT TCT GAT GTT CG |
| CCL-17 | ACC ACG TCT TCA GCT TTC TAA G | TTC TCT GCA GCA CAT CCA C |
